# CTCF orchestrates long-range cohesin-driven V(D)J recombinational scanning

**DOI:** 10.1101/2020.01.01.891473

**Authors:** Zhaoqing Ba, Jiangman Lou, Edward W. Dring, Adam Yongxin Ye, Sherry G. Lin, Suvi Jain, Kyong-Rim Kieffer-Kwon, Rafael Casellas, Frederick W. Alt

## Abstract

RAG endonuclease initiates V(D)J recombination in progenitor (pro)-B cells^1^. Upon binding a recombination center (RC)-based J_H_, RAG scans upstream chromatin via loop extrusion, potentially mediated by cohesin^2–10^, to locate Ds and assemble a DJ_H_-based RC^11^. CTCF looping factor-bound elements (CBEs) within IGCR1 upstream of Ds impede RAG-scanning^12–15^; but their inactivation allows scanning to proximal V_H_s where additional CBEs activate rearrangement and impede scanning any further upstream^15, 16^. Distal V_H_ utilization is thought to involve diffusional RC access following large-scale *Igh* locus contraction^17–23^. Here, we test the potential of linear RAG-scanning to mediate distal V_H_ usage in G1-arrested, *v-Abl-*pro-B cell lines^24, 25^, which undergo robust D-to-J_H_ but little V_H_-to-DJ_H_ rearrangements, presumably due to lack of locus contraction^11, 15^. Through an auxin-inducible approach^26, 27^, we degrade the cohesin-component Rad21^4, 7, 27^ or CTCF^7, 9^ in these G1-arrested lines, which maintain substantial viability throughout four-day experiments. Rad21 degradation eliminated all V(D)J recombination and RAG-scanning-associated interactions, except RC-located DQ52-to-J_H_ joining in which synapsis occurs by diffusion^11^. Remarkably, while CTCF degradation suppressed most CBE-based chromatin interactions, it promoted robust RC interactions with, and robust V_H_-to-DJ_H_ joining of, distal V_H_s, with patterns similar to those of “locus-contracted” primary pro-B cells. Thus, down-modulation of CTCF-bound scanning-impediment activity promotes cohesin-driven RAG-scanning across the 2.7Mb *Igh* locus.

---

Cohesin is a highly conserved chromosome-associated multi-subunit ring-shaped ATPase complex built upon subunits belonging to the structural maintenance of chromosomes (Smc) family and is important for diverse chromosome-based processes^28–33^. The cohesin complex is proposed to form contact loop domains by extrusion of chromatin until reaching convergent CTCF-bound CBE anchors^2–10^. In this regard, cohesin extrudes both naked and nucleosome-containing DNA in an NIPBL-MAU2 and ATP-hydrolysis-dependent manner *in vitro*^34, 35^. Our studies implicated the cohesin complex in chromatin loop extrusion-mediated mechanisms of lymphocyte V(D)J and IgH class switch recombination^11, 14–16, 36, 37^. To further elucidate potential cohesin role(s) in chromatin scanning during long-range V(D)J recombination, we employed a mini auxin-inducible degron (mAID) approach^4, 7, 26, 27^ to conditionally degrade Rad21, a core component of the cohesin complex, in *v-Abl*-kinase-transformed, *Eμ-Bcl2*-expressing mouse pro-B cells (“*v-Abl* pro-B cells”). These cells can be viably arrested in the G1 cell-cycle stage by treatment with Abl kinase inhibitor (STI-571), activating RAG1/2 endonuclease and V(D)J recombination^24, 25^. We targeted the last codon of both mouse *Rad21* alleles to introduce in-frame sequences that encode mAID and a green fluorescent protein (GFP) marker in a RAG2-deficient *v-Abl* pro-B line (Extended Data Fig. 1a,c). We then targeted a transgene into the *Rosa26* locus that over-expresses the OsTir1-V5 protein that binds mAID in the presence of auxin (Indole-3-acetic acid, IAA) to trigger proteasome-dependent degradation of the mAID-fused Rad21 protein (Extended Data Fig. 1a-d). The resulting RAG2-deficient line is referred to as the “Rad21-degron *v-Abl* pro-B line”.

To test effects of cohesin depletion on *Igh* loop domain formation, we split Rad21-degron *v-Abl* cells into two sub-populations, one non-treated (“NT”) and the other treated with IAA for 6 hours and then with STI-571 for 4 days to induce G1 arrest (Extended Data Fig. 1b,f). Treatment of Rad21-degron *v-Abl* cells with IAA led to depletion of mAID-GFP-tagged Rad21 to background levels by 6 hours, and ChIP-seq confirmed nearly complete depletion of chromatin-bound Rad21 at known cohesin-occupancy sites across *Igh*^11, 15, 17, 38, 39^ (Fig. 1a,b, Extended Data Fig. 2a). While there was cell loss in the initial day while cells were cycling, cohesin-depleted Rad21-degron *v-Abl* pro-B cells underwent normal STI-571-induced G1 arrest (Extended Data Fig. 1e,g), and showed no decrease in viability throughout the 4-day experimental period (Extended Data Fig. 1e). We applied the sensitive chromatin interaction 3C-HTGTS assay^15^ to G1-arrested cells at day 4, and found that cohesin loss essentially abrogated *Igh* chromatin loop domains mediated by the proximal V_H_81X-CBE^15^ or by *Igh* intronic enhancer (iEμ)/RC^11, 15^, with the exception of interaction of RC/iEμ with the closely associated, diffusion-accessible locale that harbors the infrequently rearranged (due to its poor RSS) DST4.1/D_H_3-2^11^ (Figure1c, Extended Data Fig. 2b,c). Similar to mutational inactivation of the V_H_81X-CBE^15^, Rad21 degradation completely eliminated proximal V_H_81X interaction with downstream *Igh* sequences, including the non-CBE-based RC.

**Fig. 1.**
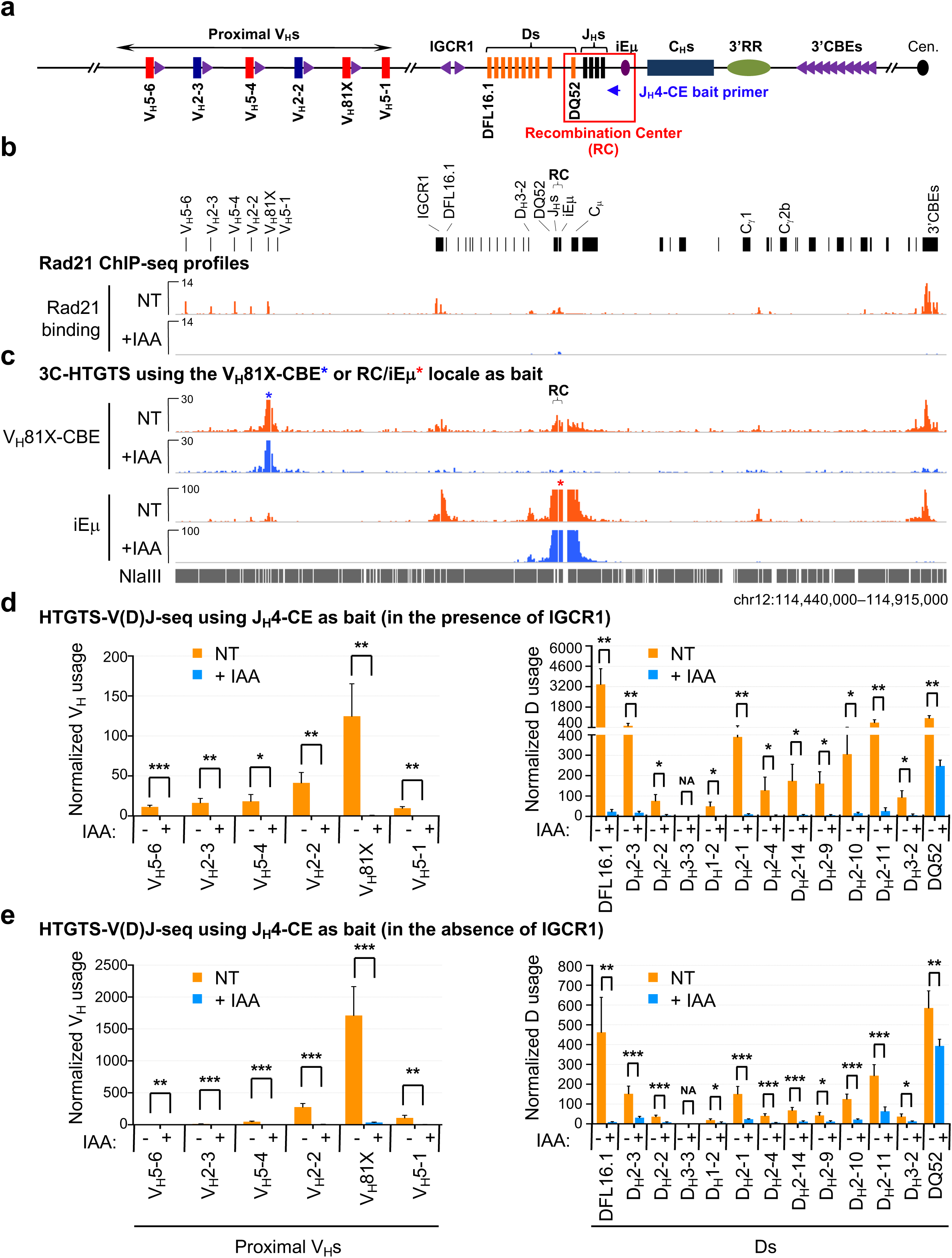
Cohesin depletion abrogates *Igh* loop domains and V(D)J rearrangements except RC-based DQ52-to-J_H_ joining in G1-arrested Rad21-degron *v-Abl* pro-B cells. **a,** Schematic of the murine *Igh* locus showing proximal V_H_s, Ds, J_H_s, C_H_s, and regulatory elements as indicated, with *Igh* recombination center (RC) that comprises J_H_-proximal DQ52 segment, four J_H_ segments, and the intronic enhancer (iEμ) highlighted. Purple arrowheads indicate organization of CBEs. Cen. centromere. Blue arrow denotes the J_H_4 coding end (CE) bait primer used for generating HTGTS-V(D)J-seq libraries. **b, c,** Representative profiles of Rad21 ChIP-seq (**b**) and 3C-HTGTS chromatin interactions using V_H_81X-CBE (blue asterisk) and RC/iEμ (red asterisk) as bait (**c**) at indicated *Igh* locus in G1-arrested Rad21-degron *v-Abl* pro-B cells without (NT) or with (+IAA) IAA treatment. See related Extended Data Fig. 2 for additional biological repeats. **d, e,** Average utilization frequencies ± s.d. of indicated proximal V_H_ (left) and D (right) segments on IGCR1-intact (**d**, *n*=3) and IGCR1-deleted (**e**, *n*=4) alleles in RAG2-complemented, G1-arrested Rad21-degron cells without (NT) or with (+IAA) IAA treatment. A J_H_4-CE bait primer was used for generating HTGTS-V(D)J-seq libraries. P values were calculated using unpaired two-tailed Student’s *t*-test, NS: P > 0.05, *: P ≤ 0.05, **: P ≤ 0.01, and ***: P ≤ 0.001. NA: not applicable. See Methods for more details.

To test effects on *Igh* V(D)J recombination, we introduced RAG2 into Rad21-degron *v-Abl* pro-B cells, with or without IAA treatment to induce mAID-GFP-tagged Rad21 depletion, and treated both with STI-571 to induce G1 arrest and activation of V(D)J recombination (Extended Data Fig.1f). After four days, we applied HTGTS-V(D)J-seq^11, 40, 41^ to NT and treated cells using a J_H_4 primer as bait^15^ to analyze both D-to-J_H_ and V_H_-to-DJ_H_ junctions (Fig. 1a,d). Similar to V(D)J joining patterns in other *v-Abl* lines, the NT Rad21-degron *v-Abl* pro-B cells displayed robust D-to-J_H_ rearrangements, with the J_H_-distal DFL16.1 having highest rearrangement frequency and J_H_-proximal DQ52 an intermediate frequency^11^ (Fig. 1d, right panel, Supplementary Table 1). We also observed low-level rearrangement of the most proximal V_H_s and little or no rearrangement of distal V_H_s^11, 14, 15^ (Fig.1d, left panel, Supplementary Table 1). Strikingly, Rad21 depletion abolished all V_H_ rearrangements and largely eliminated all D rearrangements, except that of the J_H_-proximal DQ52, which retained substantial rearrangement levels (Fig. 1d, Supplementary Table 1). In additional studies, we found Rad21 degradation abrogated robust proximal V_H_ utilization on IGCR1-deleted alleles along with nearly complete abrogation of all D rearrangements except, again, that of proximal DQ52, which occurred at nearly 70% of control levels (Fig. 1e, Supplementary Table 1). Consistent with impeded loop extrusion, Rad21 degradation also abrogated the greatly robust *Igh* RC interaction with proximal V_H_s in the absence of IGCR1, but had little effect on RC interactions with the diffusion-accessible D_H_3-2 locale (Extended Data Fig. 2d).

The substantial level of residual DQ52 rearrangement in the various RAG-sufficient, Rad21-depleted *v-Abl* pro-B lines provides an internal control for maintenance of RAG activity and integrity of a J_H_-based RC following cohesin depletion, as DQ52 is known to substantially access RC-bound RAG via diffusion due to its RC-based location^11^. To further confirm integrity of the *Igh* RC, we performed GRO-seq on G1-arrested Rad21-degron cells, which confirmed robust sense and anti-sense transcription through the RC before and after cohesin degradation (Extended Data Fig. 3a). Furthermore, while Rad21-depletion led to expected subtle genome-wide transcriptional changes^4^ (Extended Data Fig. 3b), we detected no substantial differences in transcription levels of several genes involved in V(D)J recombination *per se* or in chromosomal regulation of V(D)J recombination (Examples highlighted in Extended Data Fig. 3b). Taken together, our findings strongly implicate cohesin-mediated loop extrusion in moving chromatin past RC-bound RAG for scanning during *Igh* V(D)J recombination.

How the 100-plus V_H_s embedded across the 2.4Mb V_H_ domain gain access to the DJ_H_ RC has been a major question. In this regard, there are over 100 CBEs across this domain in convergent orientation with the upstream IGCR1 CBE and 10 CBEs (“3’CBEs”) downstream of *Igh*^12, 13, 17, 21, 22, 42^. While most of these CBEs are not physically adjacent to V_H_s, several dozen D-proximal V_H_s have CBEs directly downstream that promote associated V_H_ utilization through interactions with the DJ_H_ RC during RAG scanning^15, 16^. In addition to IGCR1 CBEs, these proximal V_H_ CBEs are barriers to direct RAG scanning to more distal V_H_s^15^. Distal V_H_ utilization might circumvent CBE linear scanning barriers through a physical locus contraction process that brings all V_H_s into proximity with the DJ_H_ RC for diffusional access^17–23^. On the other hand, extrusion-mediated RAG scanning could, in theory, also contribute to distal V_H_ utilization if CBE impediments were neutralized or circumvented^15, 16^. To test the latter hypothesis, we asked whether we could activate RAG scanning to distal V_H_s in G1-arrested *v-Abl* pro-B cells by down-regulating CTCF scanning barriers across the locus. For this purpose, we generated a CTCF degron line from the parental RAG2-deficient *v-Abl* line using a similar approach to that used for establishing the Rad21-degron line (Extended Data Fig. 4a-e). We then followed the same experimental protocol used to study the Rad21-degron line.

Treatment of CTCF-degron *v-Abl* cells with IAA rapidly led to depletion of mAID-GFP-tagged CTCF to background levels by 6 hours, which were maintained throughout the course of the subsequent 4-day experiment (Extended data Figure 4b). Treatment with STI-571 induced normal G1 arrest of both non-treated and treated CTCF-degron *v-Abl* cells (Extended Data Fig. 4f). However, CTCF degradation moderately impacted viability of treated G1-arrested, CTCF-degron *v-Abl* cells, which decreased to approximately 55% of NT or parental control cells over the 4-day experimental course (Extended Data Fig. 4g). We also found that the non-treated CTCF-degron *v-Abl* cells had a modestly leaky phenotype for CTCF degradation^7, 9^, which provided additional insights (Extended Data Fig. 5 and legend). However, to simplify presentation of our findings, we focus here on description of the striking results obtained from comparison of the parental *v-Abl* cells and IAA-treated CTCF-degron *v-Abl* lines derived from them.

To characterize effects of CTCF degradation, we performed ChIP-seq for CTCF- and Rad21-binding across the *Igh* of G1-arrested parental and treated CTCF-degron *v-Abl* cells (Fig. 2a,b, Extended Data Fig. 6a). In parental *v-Abl* cells, there was a high density of co-localized CTCF and Rad21 peaks throughout the V_H_ locus, with more robust binding in the distal region^17, 38^. In treated CTCF-degron *v-Abl* cells, CTCF occupancy across *Igh* was greatly diminished; but residual binding^9, 17^ was retained at certain CBEs in the distal region (e.g. V_H_1-67, V_H_1-11), as well as CBEs associated with proximal V_H_5-6, V_H_2-3, and CBE1 of IGCR1 (Fig. 2b, Extended Data Fig. 6a). Notably, CTCF binding increased at the RC upon CTCF depletion (Fig. 2b, Extended Data Fig. 6a). While increases in apparent CTCF-binding to the non-CBE-containing RC upon CTCF depletion could be considered surprising, it may likely occur indirectly due to cohesin-mediated loop extrusion-driven dynamic associations of the RC sub-loop anchor with residual-CTCF-bound CBEs across *Igh.* The, apparently, uneven depletion of CTCF at various CBEs may reflect differential intrinsic CTCF-binding affinity, local chromatin context and/or other factors^43–45^. Upon CTCF depletion, Rad21 occupancy throughout *Igh* was also greatly diminished, with residual binding sites and levels largely reflecting those of CTCF (Fig. 2b, Extended Data Fig. 6a). Residual CTCF binding upon CTCF depletion likely reflects average states of depletion of different cells in the population of G1-arrested CTCF-degron *v-Abl* cells, with the latter also being influenced by loss of particular cells in which CTCF depletion reaches a critical level for viability. Therefore, occurrence of continuous sub-populations of G1-arrested, CTCF-degron *v-Abl* cells in which residual CTCF maintains viability implies that such cells may also maintain RAG-scanning in the face of variable CTCF depletion across *Igh.* If so, this would further predict increased RC scanning anchor interactions with the distal V_H_ locus and increased utilization of distal V_H_s.

**Fig. 2.**
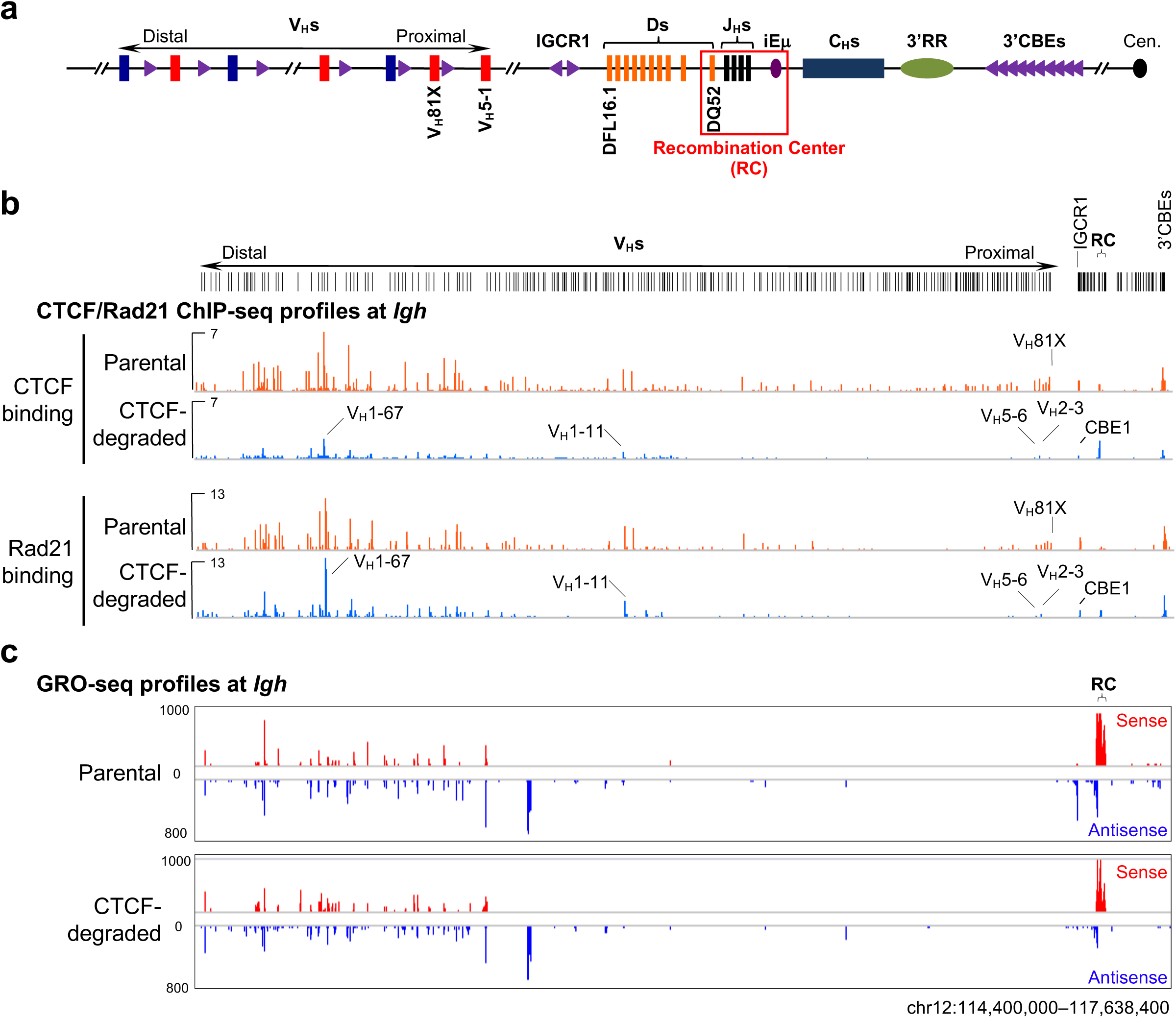
Effects of CTCF degradation on chromatin CTCF/Rad21-binding and transcription across *Igh* locus in G1-arrested CTCF-degron *v-Abl* pro-B cells. **a,** Schematic of the entire murine *Igh* locus with details as shown in Fig. 1a. **b, c,** Representative profiles of ChIP-seq (**b**) using CTCF (top panels) and Rad21 (bottom panels) antibodies and GRO-seq (**c**) across the entire *Igh* locus as indicated in G1-arrested parental and CTCF-degraded *v-Abl* pro-B cells. CBEs with residual CTCF-binding are indicated in **b**. The entire V_H_ locus is diagrammed at the top. Additional biological repeats of each experiment are shown in related Extended Data Fig. 6. See Methods for more details.

GRO-seq confirmed that transcription of the RC and distal V_H_s occurred similarly in parental and treated CTCF-degron *v-Abl* lines (Fig. 2c, Extended Data Fig.6b), predicting an intact RC and accessibility of distal V_H_s if distal scanning reached them. Correspondingly, IAA-treated, G1-arrested CTCF-degron *v-Abl* lines had a dramatic increase in utilization of many V_H_s upstream of proximal V_H_81X and V_H_2-2, including transcribed distal V_H_s that lie several Mb upstream (Fig. 3a-c, Supplementary Table 2). Remarkably, the patterns of V_H_ rearrangements in CTCF-degraded *v-Abl* cells were reminiscent of those in “locus-contracted” bone marrow (BM) pro-B cells in terms of similar highly rearranging V_H_ clusters and comparable levels of some distal V_H_ rearrangements (Fig. 3c,d, Supplementary Table 2). Moreover, compared to parental *v-Abl* cells, the V_H_DJ_H_ to DJ_H_ ratio in CTCF-degraded *v-Abl* cells reached that of BM pro-B cells, confirming that CTCF deletion greatly increased the overall rate of V_H_ to DJ_H_ joining by permitting relatively unimpeded far upstream scanning for diverse V_H_ rearrangements to DJ_H_ RCs (Fig. 3). Utilization of the most proximal functional V_H_s (V_H_81X and V_H_2-2) decreased relative to that of IGCR1-inactivated normal G1-arrested *v-Abl* cells or even that of normal pro-B cells. This latter finding may reflect CTCF-depletion dampening at V_H_81X- and V_H_2-2-associated CBEs, as well as at IGCR1 CBEs (Fig. 2b, Extended Data Fig. 6a). Taken together, these results confirm the hypothesis that accessibility of distal V_H_s to RC-based RAG is greatly enhanced by broad diminution of CTCF-based scanning impediments across *Igh*.

**Fig. 3.**
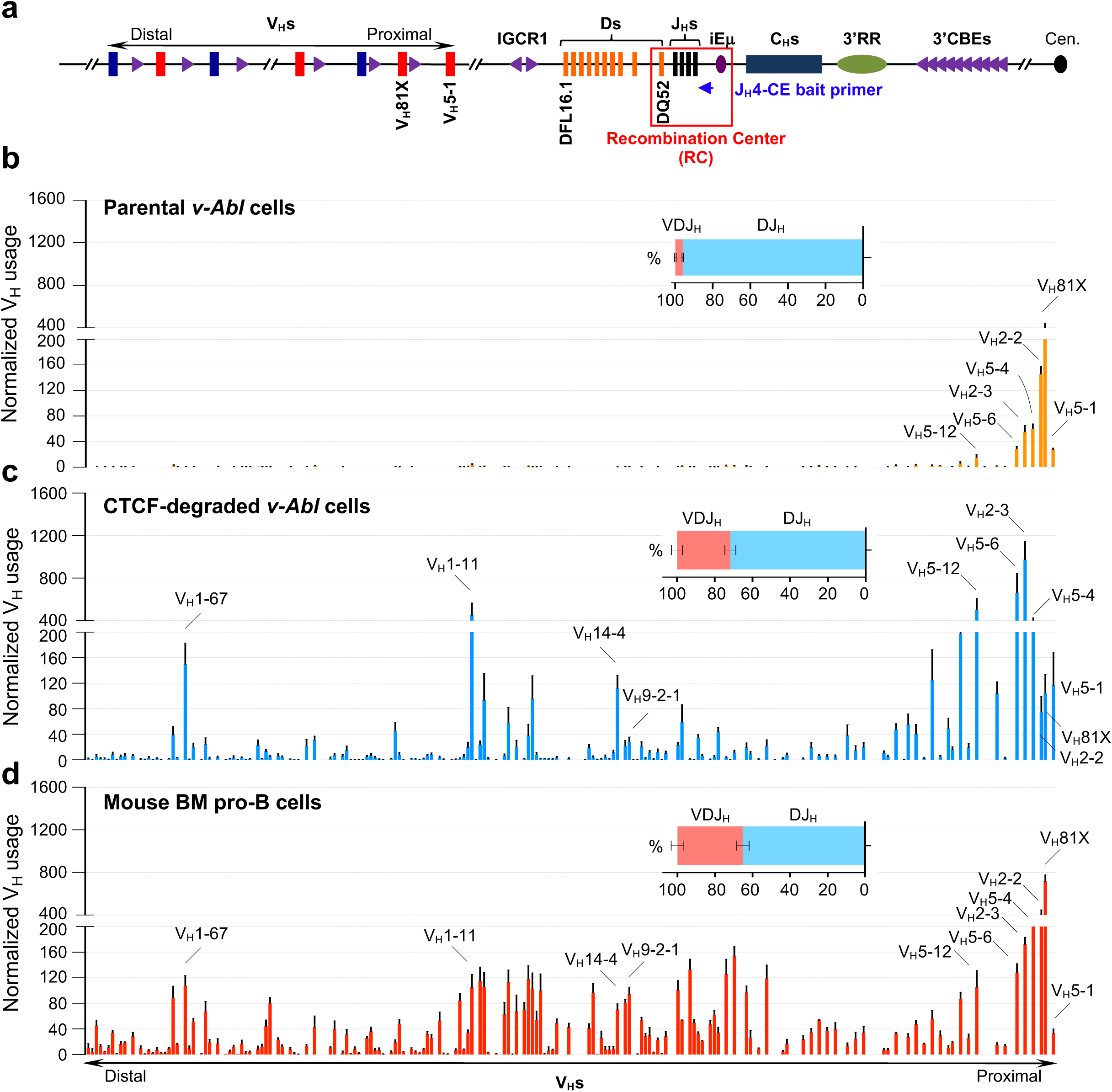
CTCF degradation activates distal V_H_ rearrangements in G1-arrested CTCF-degron *v-Abl* pro-B cells. **a,** Schematic of the murine *Igh* locus. Blue arrow denotes the J_H_4-CE bait primer used for generating HTGTS-V(D)J-seq libraries. Other details are as in Fig. 2a. **b-d,** Average utilization frequencies ± s.d. of all V_H_ segments in RAG2-complemented, G1-arrested parental (**b**, *n*=4), CTCF-degraded (**c**, *n*=6) *v-Abl* pro-B cells and mouse BM pro-B cells (**d**, *n*=3). Average percentage ± s.d. of VDJ_H_ and DJ_H_ rearrangements among total V(D)J junctions for each sample are indicated. See Methods for more details. For comparison of V_H_ utilization patterns among samples, the location of several V_H_s are indicated.

3C-HTGTS from an RC/iEμ bait on G1-arrested parental *v-Abl* cells confirmed lack of interaction with the more distal V_H_s (Fig. 4a, b, top panel). Remarkably, in treated CTCF-degron *v-Abl* cells, the RC/iEμ gained robust interactions with sequences across the entire V_H_ locus, including the most distal V_H_s, with patterns reminiscent of those in normal locus-contracted BM pro-B cells (Fig. 4b, middle and bottom panels). We confirmed these chromatin interaction analyses with other RC-based baits (Extended Data Fig. 7a-c). Notably, many V_H_s near prominent RC/iEμ-interacting peaks corresponded to those with greatly enhanced utilization (Fig. 3c,4b). In addition, we observed that, compared to parental *v-Abl* cells, the RC retained interactions with IGCR1, had diminished interactions with V_H_81X- and V_H_2-2-associated CBEs, and had increased interactions with proximal V_H_2-3 and V_H_5-6 in CTCF-degraded *v-Abl* cells (Fig. 4b, Extended Data Fig. 7b,c, top two panels). These chromatin interaction changes corresponded well to proximal V_H_ utilization changes (Fig. 3b,c). Together with our finding of CTCF degradation-dependent enhancement of distal V_H_ rearrangements, our chromatin interaction findings indicate that diminution of CTCF-based scanning impediments allows linear interaction by the RC and, correspondingly, RC-bound RAG, to proceed across the entire 2.7Mb *Igh* V_H_ locus, likely to different distances in individual cells in the population.

**Fig. 4.**
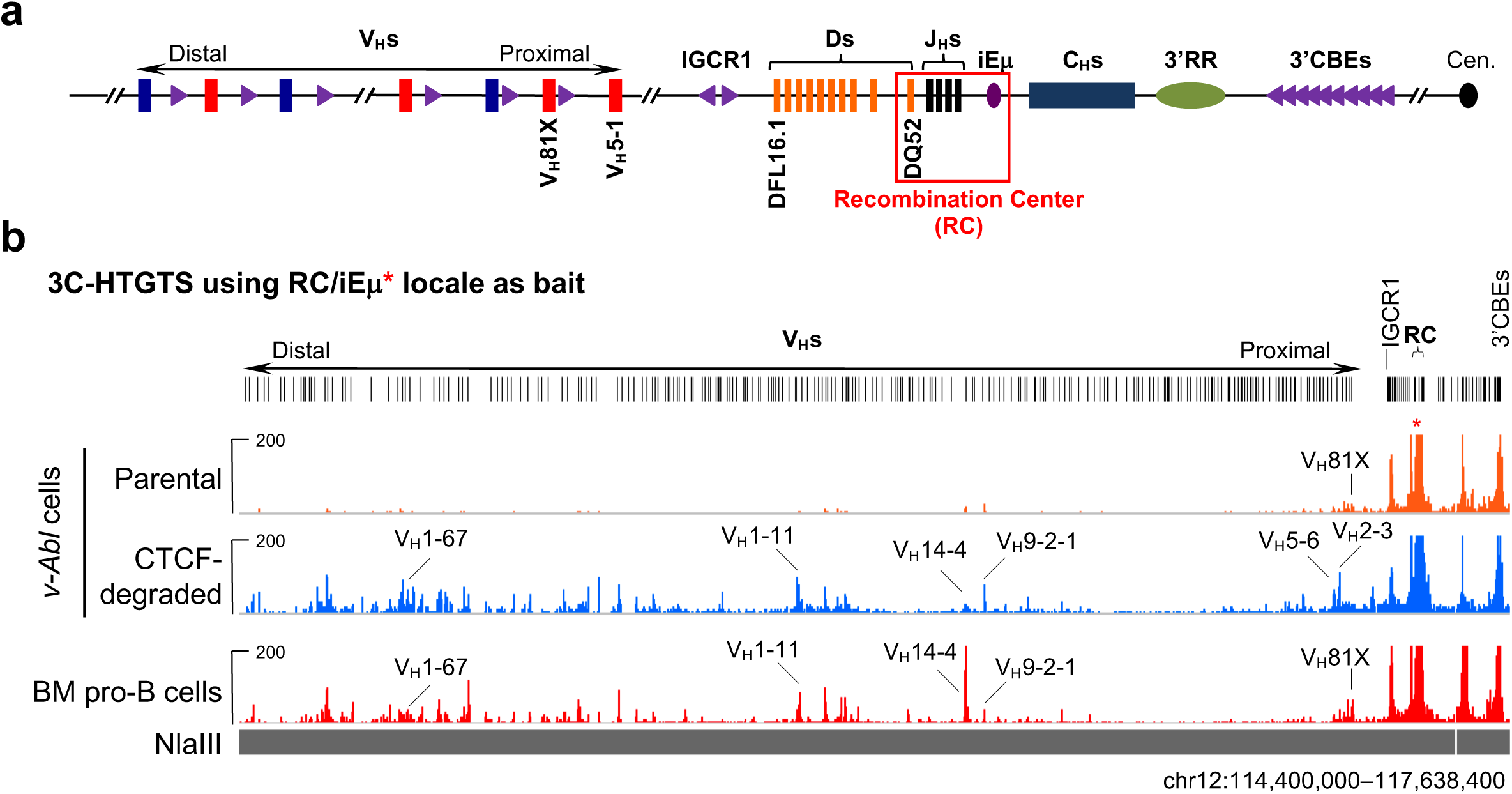
CTCF degradation restores long-distance *Igh* RC interactions with distal V_H_ sequences in G1-arrested CTCF-degron *v-Abl* pro-B cells. **a,** Schematic of the entire murine *Igh* locus with details as shown in Fig. 2a. **b,** Representative 3C-HTGTS chromatin interaction profiles of RC/iEμ bait (red asterisk) across the entire *Igh* locus in G1-arrested parental (top), CTCF-degraded (middle) *v-Abl* pro-B cells and RAG2-deficient mouse BM pro-B cells (bottom). See Methods for more details. For comparison, several distal interaction peaks located proximal to V_H_s highlighted in Fig. 3 are indicated. The entire V_H_ locus is diagrammed at the top. See related Extended Data Fig. 7 for additional 3C-HTGTS profiles with other RC-based baits.

Our cohesin-depletion studies in G1-arrested *v-Abl* pro-B cells strongly support cohesin-mediated loop extrusion as a major driver in moving chromatin past RC-bound RAG for *Igh* V(D)J recombinational scanning. Moreover, our studies show that RAG scanning can progress linearly across the entire V_H_ locus upon down-modulation of CBE impediment activity allowing other local scanning impediments (e.g. transcription, Fig. 2c, Extended Data Fig.6b)^11, 37, 46^ to mediate increased access to particular sets of V_H_s (Fig. 4, Extended Data Fig. 7,8 and legend). Indeed, CTCF-degradation actually increased the overall level of V_H_-to-DJ_H_ rearrangements, including those of distal V_H_s, in *v-Abl* pro-B lines to levels similar to those found in primary pro-B cells (Fig. 3). While CTCF depletion provides proof of principle that linear RAG scanning can access to distal V_H_s, physiological modulation of scanning activity may involve mechanisms that down-regulate CBE impediments and/or mechanisms that circumvent impediments by modulating cohesin activity. One such possibility would be modulating activity of the WAPL cohesin unloader, given that WAPL depletion extends length of CBE-based loops^5, 7^. Thus, mechanisms that, directly or indirectly, impact CTCF-bound CBE scanning impediments during early B cell development could contribute to IgH variable region repertoire diversification via linear scanning. Cohesin-mediated loop extrusion may be similarly regulated in the context of V(D)J recombination in other antigen receptor loci including Igκ^47, 48^ and T cell receptor loci^36, 49, 50^, consistent with cohesin being required for *TCRa* rearrangement in developing T cells^49^. The system we describe for analyzing cohesin-complex protein functions in G1-arrested *v-Abl* pro-B cells should be useful for further testing functions and mechanisms of these or other factors in physiological chromatin loop extrusion *in vivo*.

## ACKNOWLEDGEMENTS

We thank lab members for stimulating discussions. This work was supported by NIH R01 AI020047 (to F.W.A.). R.C. is partially funded by the NIH Regulome Project. F.W.A. is an investigator of the Howard Hughes Medical Institute. Z.B. was a Cancer Research Institute Irvington fellow.

## AUTHOR CONTRIBUTIONS

Z.B., J.L., R.C., and F.W.A. designed the study; Z.B. and J.L. performed most of the experiments with assistance from E.D., S.G.L., and K.-R.K.-K. on certain experiments; A.Y.Y. designed some of the bioinformatics pipelines for data analysis; S.J. provided helpful discussions early in the study; Z.B., J.L., and F.W.A. analyzed and interpreted data, designed figures, and wrote the paper. R.C., S.G.L., and S.J. helped polish the paper. F.W.A. supervised the study.

## AUTHOR INFORMATION

The authors declare no competing financial interests. Correspondence and requests for materials should be addressed to F.W.A. F.W.A. is a co-founder of Otoro Biopharmaceuticals.

## EXTENDED DATA FIGURE LEGENDS

**Extended Data Fig. 1.**
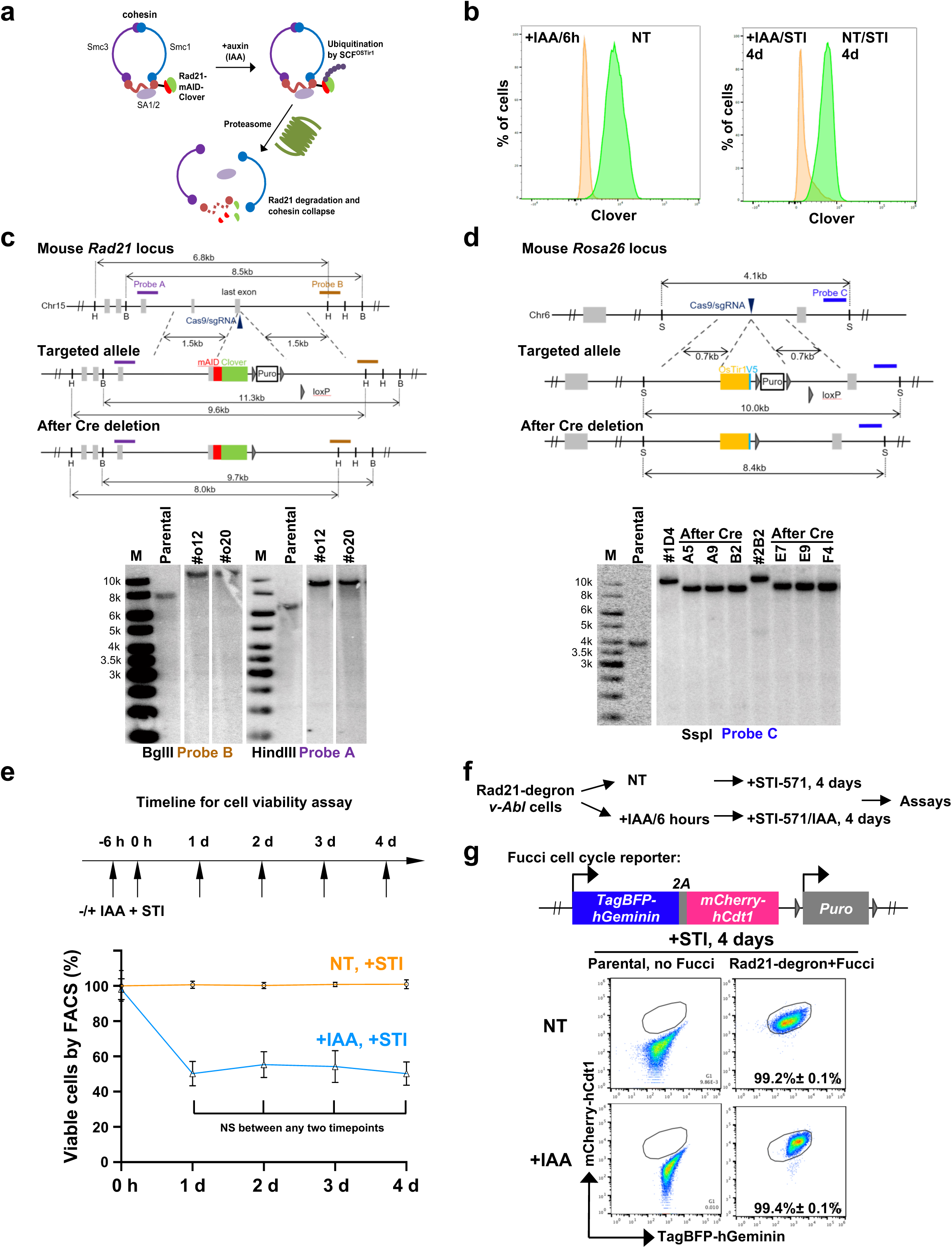
Generation and characterization of Rad21-degron *v-Abl* pro-B lines. **a,** Diagram of the Rad21-degron mouse *v-Abl* pro-B line. **b,** Representative flow-cytometry plots showing the percentage of Clover-positive Rad21-degron *v-Abl* cells that are non-treated (NT) and treated by IAA (+IAA) for 6h (left) followed by treatment with STI-571 (without or with IAA) for 4 days to induce G1 arrest (right). **c, d,** Top: Schematic of the targeting strategy for introducing in-frame mAID-Clover sequences into the last codon of mouse both endogenous *Rad21* alleles (**c**), and OsTir1-V5 expression cassette into both endogenous *Rosa26* alleles (**d**) in a *rag2^-/-^; Eμ-Bcl2^+^ v-Abl* pro-B line. Positions of Cas9/sgRNAs and Southern blot probes are indicated. H: HindIII; B: BglII; S: SspI. Bottom: Southern blot confirmation of correctly targeted alleles as indicated. **e,** Time-course cell viability assay for Rad21-degron *v-Abl* pro-B cells without (-IAA, NT) and with (+IAA) IAA treatment following STI-571 treatment for G1-arrest (+STI). Top: the assay timeline. Bottom: time-course cell viability curves. Average percentage ± s.d. of viable cells for each timepoint and under each condition was shown (*n*=4 experiments with biologically independent clones). NS: P > 0.05. **f,** Diagram of the experimental strategy with Rad21-degron *v-Abl* cells for various assays presented in this study. **g,** Fucci cell cycle assay of Rad21-degron *v-Abl* cells without (NT) and with (+IAA) IAA treatment following 4-day STI-571 treatment for G1-arrest. Representative flow-cytometry plots and average percentage ± s.d. of cells arrested in G1 stage at indicated condition (*n*=3) are shown. See Methods for details.

**Extended Data Fig. 2.**
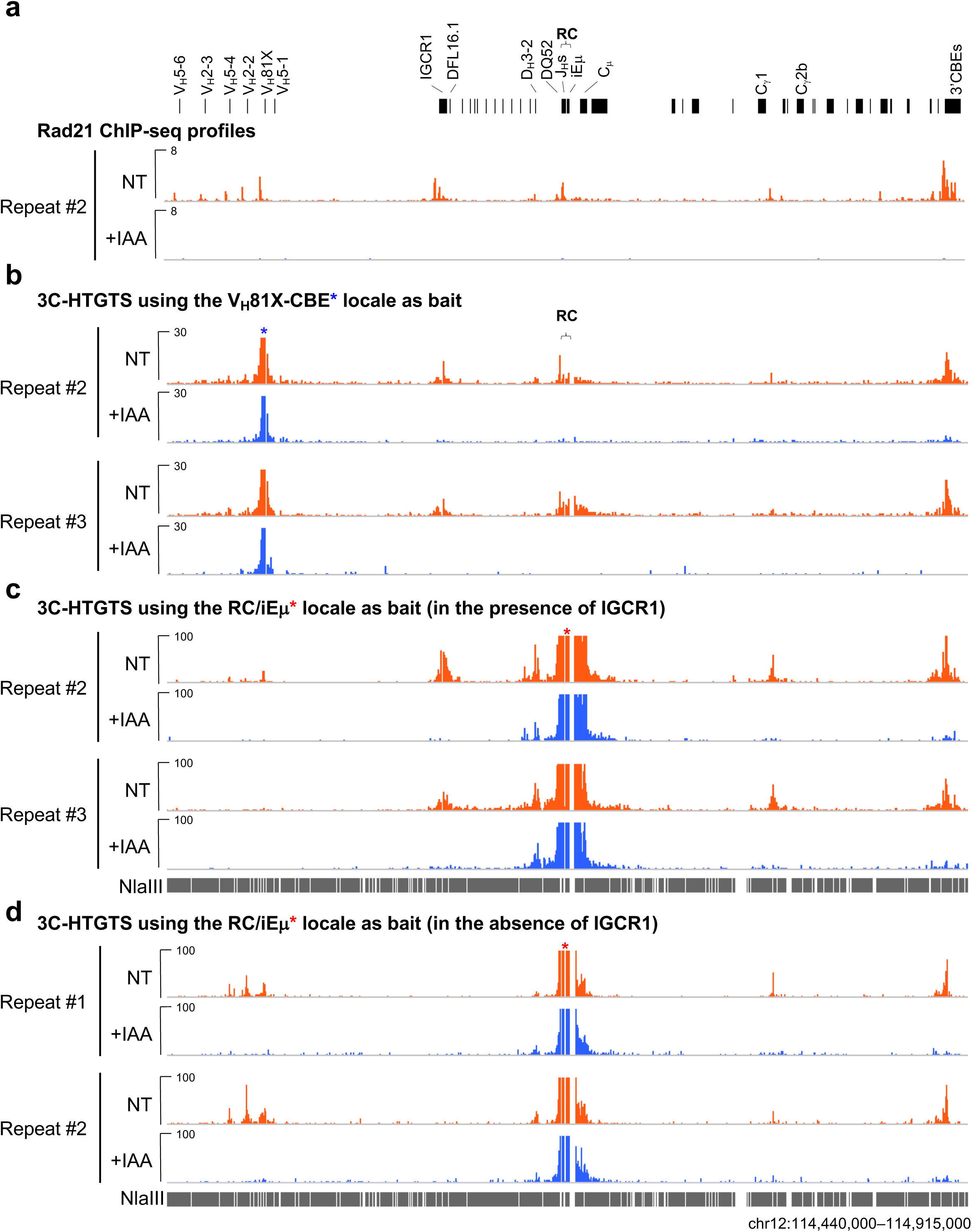
Cohesin depletion abrogates *Igh* loop domains in the presence or absence of IGCR1 in G1-arrested Rad21-degron *v-Abl* pro-B cells. **a,** An additional biologically independent repeat of Rad21 ChIP-seq profiles at indicated *Igh* locus in G1-arrested Rad21-degron *v-Abl* pro-B cells without (NT) or with (+IAA) IAA treatment as shown in Fig. 1b. **b, c,** Two additional biologically independent repeats for 3C-HTGTS chromatin interaction profiles of V_H_81X-CBE bait (**b**, blue asterisk) and iEμ bait (**c**, red asterisk) at indicated *Igh* locus in G1-arrested Rad21-degron *v-Abl* pro-B cells without (NT) or with (+IAA) IAA treatment as shown in Fig. 1c. **d,** Two biologically independent repeats for 3C-HTGTS chromatin interaction profiles of iEμ bait (red asterisk) at indicated *Igh* locus in G1-arrested IGCR1-deleted Rad21-degron *v-Abl* pro-B cells without (NT) or with (+IAA) IAA treatment. See Methods for more details.

**Extended Data Fig. 3.**
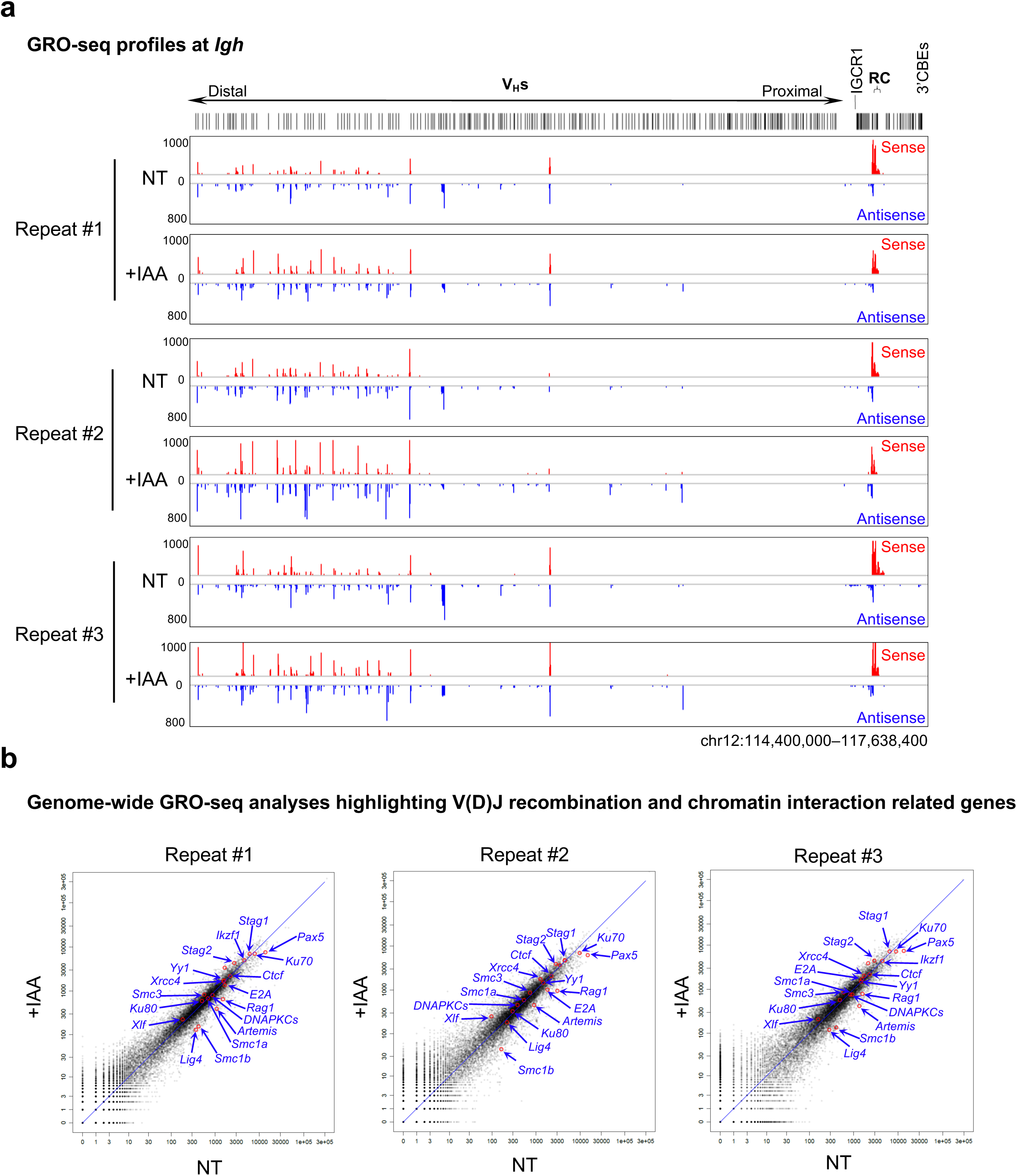
Effects of cohesin loss on *Igh* and genome-wide gene transcription in G1-arrested Rad21-degron *v-Abl* pro-B cells. **a,** Three biologically independent GRO-seq repeats of G1-arrested Rad21-degron *v-Abl* cells without (NT) or with (+IAA) IAA treatment across the entire *Igh* locus as indicated. **b,** Scatter plots of transcriptome-wide GRO-seq (three biologically independent repeats) counts in G1-arrested Rad21-degron *v-Abl* cells without (NT, *x* axis) or with (+IAA, *y* axis) IAA treatment. Representative known requisite genes for V(D)J recombination and chromatin interaction are highlighted by red circles and blue arrows in each of the three scatter plots. See Methods for more details.

**Extended Data Fig. 4.**
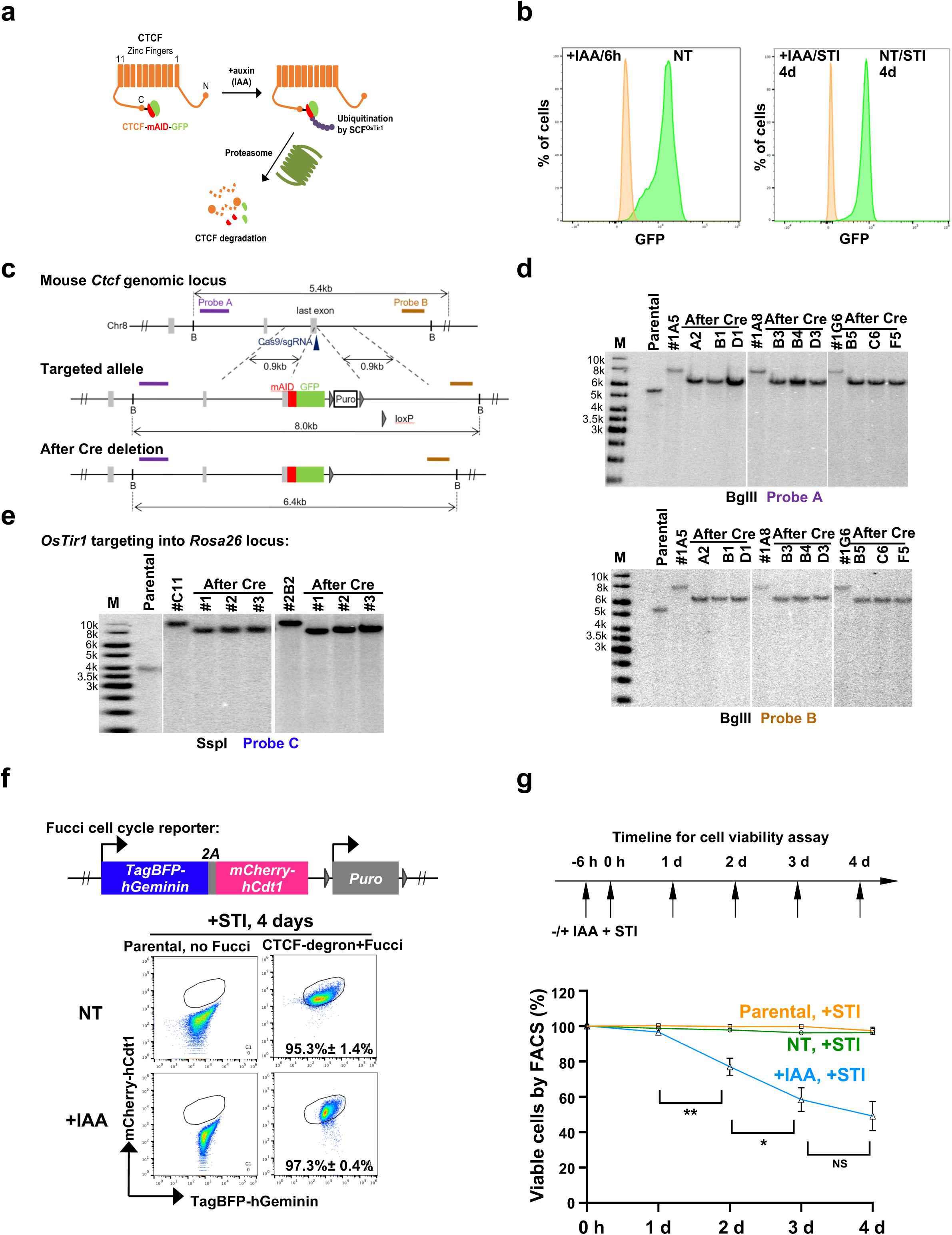
Generation and characterization of CTCF-degron *v-Abl* pro-B lines. **a,** Diagram of the CTCF-degron mouse *v-Abl* pro-B line. **b,** Representative flow-cytometry plots showing the percentage of GFP-positive CTCF-degron *v-Abl* cells that are non-treated (NT) and treated by IAA (+IAA) for 6h (left) followed by treatment with STI-571 (without or with IAA) for 4 days to induce G1 arrest (right). **c,** Schematic of the targeting strategy for introducing in-frame mAID-GFP sequences into the last codon of mouse both *Ctcf* alleles in a *rag2^-/-^; Eμ-Bcl2^+^ v-Abl* pro-B line. Positions of Cas9/sgRNAs and Southern blot probes are indicated. B: BglII. **d, e,** Southern blot confirmation of correctly targeted *Ctcf* (**d**) and *Rosa26* (**e**) alleles as indicated. Targeting OsTir1-V5 cassette into *Rosa26* alleles follows the same targeting strategy as shown in Extended Data Fig. 1d. **f,** Fucci cell cycle assay of CTCF-degron *v-Abl* cells without (NT) and with (+IAA) IAA treatment following 4-day STI-571 treatment for G1-arrest. Representative flow-cytometry plots and average percentage ± s.d. of cells arrested in G1 stage at indicated condition (*n*=3) are shown. **g,** Time-course cell viability assay for CTCF-degron *v-Abl* pro-B cells without (-IAA, NT) and with (+IAA) IAA treatment following STI-571 treatment for G1-arrest (+STI). Top: the assay timeline. Bottom: time-course cell viability curves. Average percentage ± s.d. of viable cells for each timepoint and under each condition was shown (*n*=3 experiments with biologically independent clones). NS: P > 0.05, *: P ≤ 0.05, **: P ≤ 0.01. See Methods for details.

**Extended Data Fig. 5.**
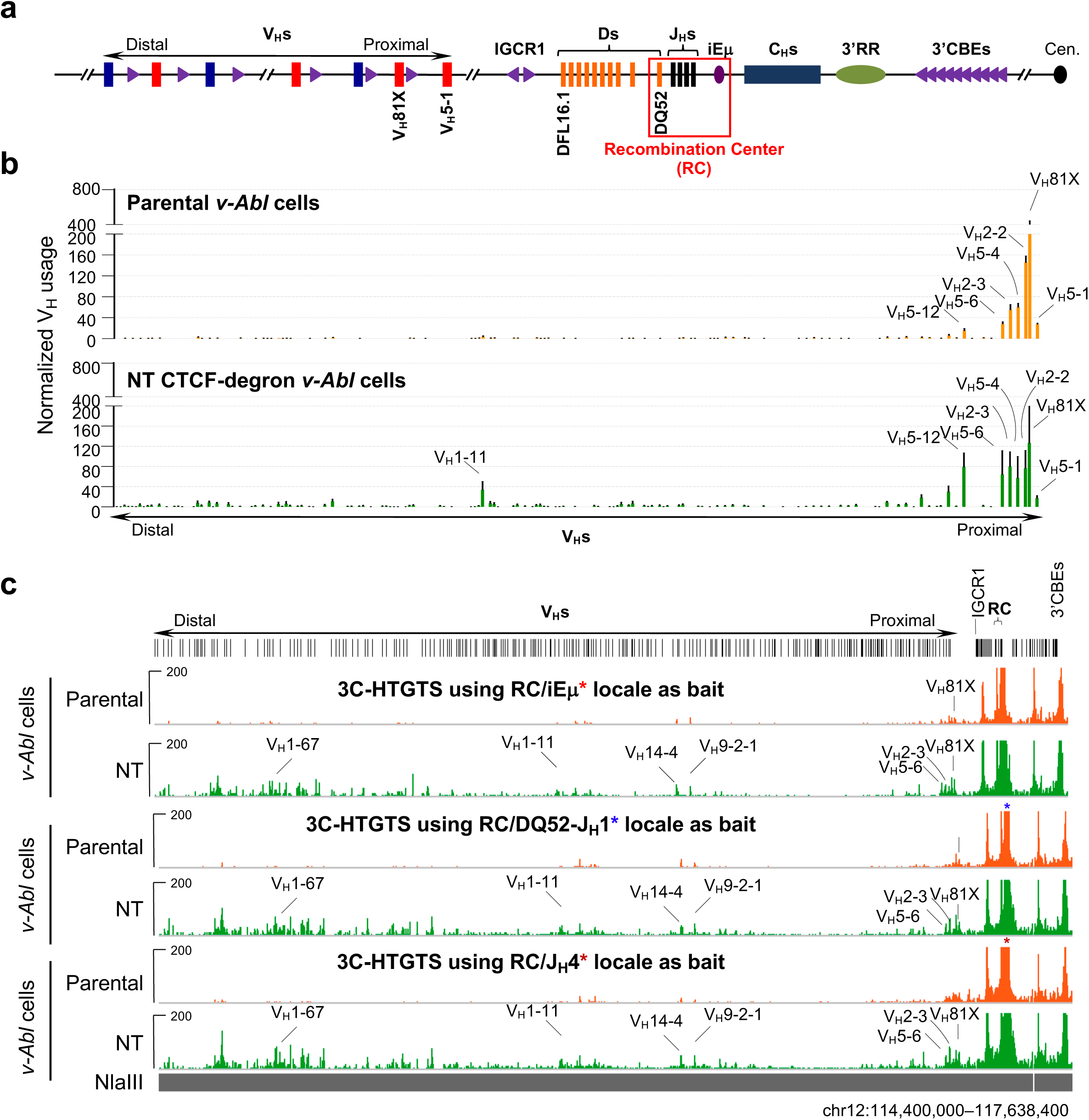
CTCF-degron *v-Abl* pro-B lines show modest leakiness. **a,** Schematic of the murine *Igh* locus as in Fig. 2a. **b**, Average utilization frequencies ± s.d. of all V_H_s in RAG2-complemented, G1-arrested parental *v-Abl* pro-B cells (top, *n*=4) and non-treated CTCF-degron *v-Abl* pro-B cells (bottom, *n*=6). For comparison, locations of several V_H_s are indicated. **c**, Representative 3C-HTGTS chromatin interaction profiles of RC/iEμ (red asterisk), RC/DQ52-J_H_1 (blue asterisk), and RC/J_H_4 (purple asterisk) baits across the *Igh* locus in G1-arrested parental *v-Abl* pro-B cells (top) and non-treated CTCF-degron *v-Abl* pro-B cells (bottom). See Methods for details. For comparison, several distal interaction peaks near V_H_s highlighted in Fig.3 are indicated. The V_H_ locus is diagrammed at the top. See Extended Data Fig. 7a for diagrams of 3C-HTGTS RC baits employed. To facilitate direct comparisons, the HTGTS-V(D)J-seq and 3C-HTGTS data for the Parental *v-Abl* cells are the same as those presented in figED. 3, 4 and Extended Data Fig. 7. **Discussion:** The apparent greater leakiness with respect to RC interactions versus RAG scanning activity across the upstream locus might result from the latter being done in RAG2-sufficient cells. As proposed to explain a related phenomenon involving dCas9-binding, extrusion of chromatin impediments past the RC may be more efficient when not RAG-bound^11^.

**Extended Data Fig. 6.**
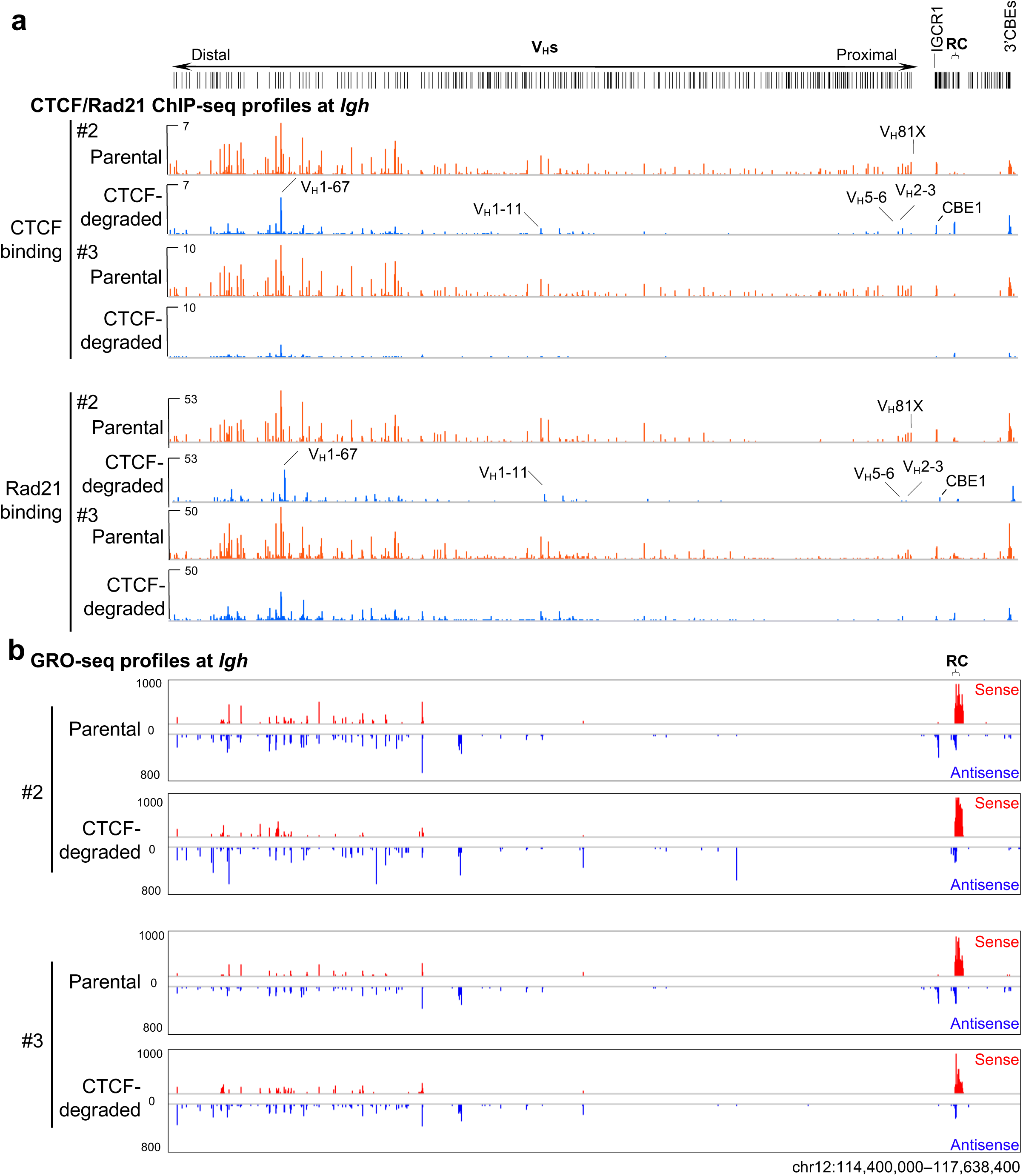
Effects of CTCF degradation on chromatin CTCF/Rad21-binding and transcription across *Igh* locus in G1-arrested CTCF-degron *v-Abl* pro-B cells. **a, b,** Two additional biologically independent repeats of ChIP-seq (**a**) using CTCF (top panels) and Rad21 (bottom panels) antibodies and GRO-seq (**b**) across the entire *Igh* locus as indicated in G1-arrested parental and CTCF-degraded *v-Abl* pro-B cells. CBEs with residual CTCF-binding are indicated in **a**. The entire V_H_ locus is diagrammed at the top. See Methods for more details.

**Extended Data Fig. 7.**
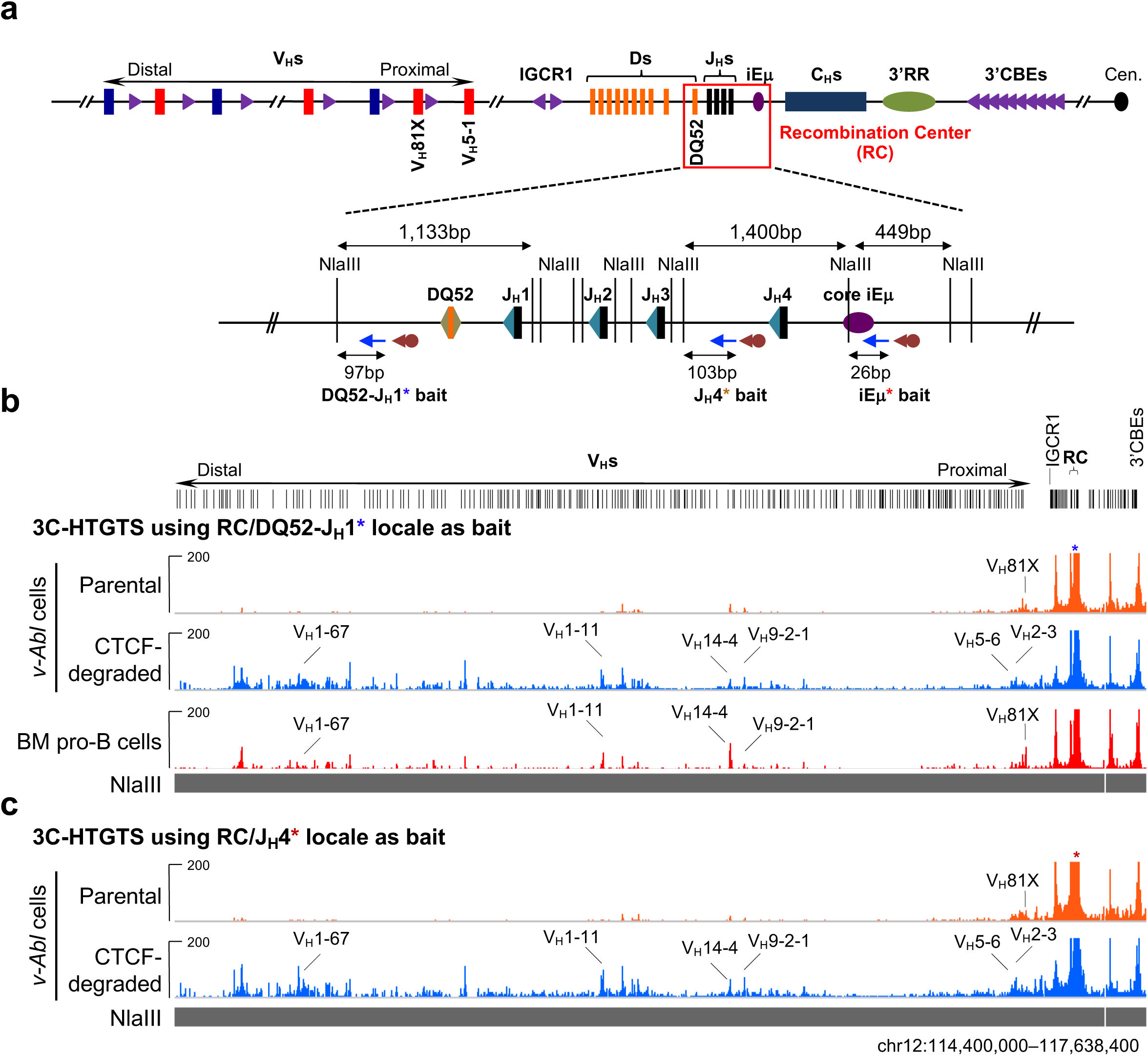
CTCF degradation restores long-distance *Igh* RC interactions with distal V_H_ sequences in G1-arrested CTCF-degron *v-Abl* pro-B cells. **a,** Top: Schematic of the murine *Igh* locus as in Fig. 2a. Bottom: Zoom-in schematic of *Igh* RC showing 3C-HTGTS assay baiting strategies used for iEμ (red asterisk), DQ52-J_H_1 (blue asterisk), and J_H_4 (purple asterisk) locales. NlaIII restriction fragments and the relative positions of biotinylated (cayenne arrow) and nested (blue arrow) PCR primers employed are illustrated. Brown and teal blue triangles represent DQ52-12RSSs and J_H_-23RSSs, respectively. **b,** Representative 3C-HTGTS chromatin interaction profiles of RC/DQ52-J_H_1 (blue asterisk) across the entire *Igh* locus in G1-arrested parental (top), CTCF-degraded (middle) *v-Abl* pro-B cells and RAG2-deficient mouse BM pro-B cells (bottom). **c,** Representative 3C-HTGTS chromatin interaction profiles of RC/J_H_4 (purple asterisk) across the entire *Igh* locus in G1-arrested parental (top) and CTCF-degraded (bottom) *v-Abl* pro-B cells. See Methods for details. For comparison, several distal interaction peaks located proximal to V_H_s highlighted in Fig. 3 are indicated. The entire V_H_ locus is diagrammed at the top.

**Extended Data Fig. 8.**
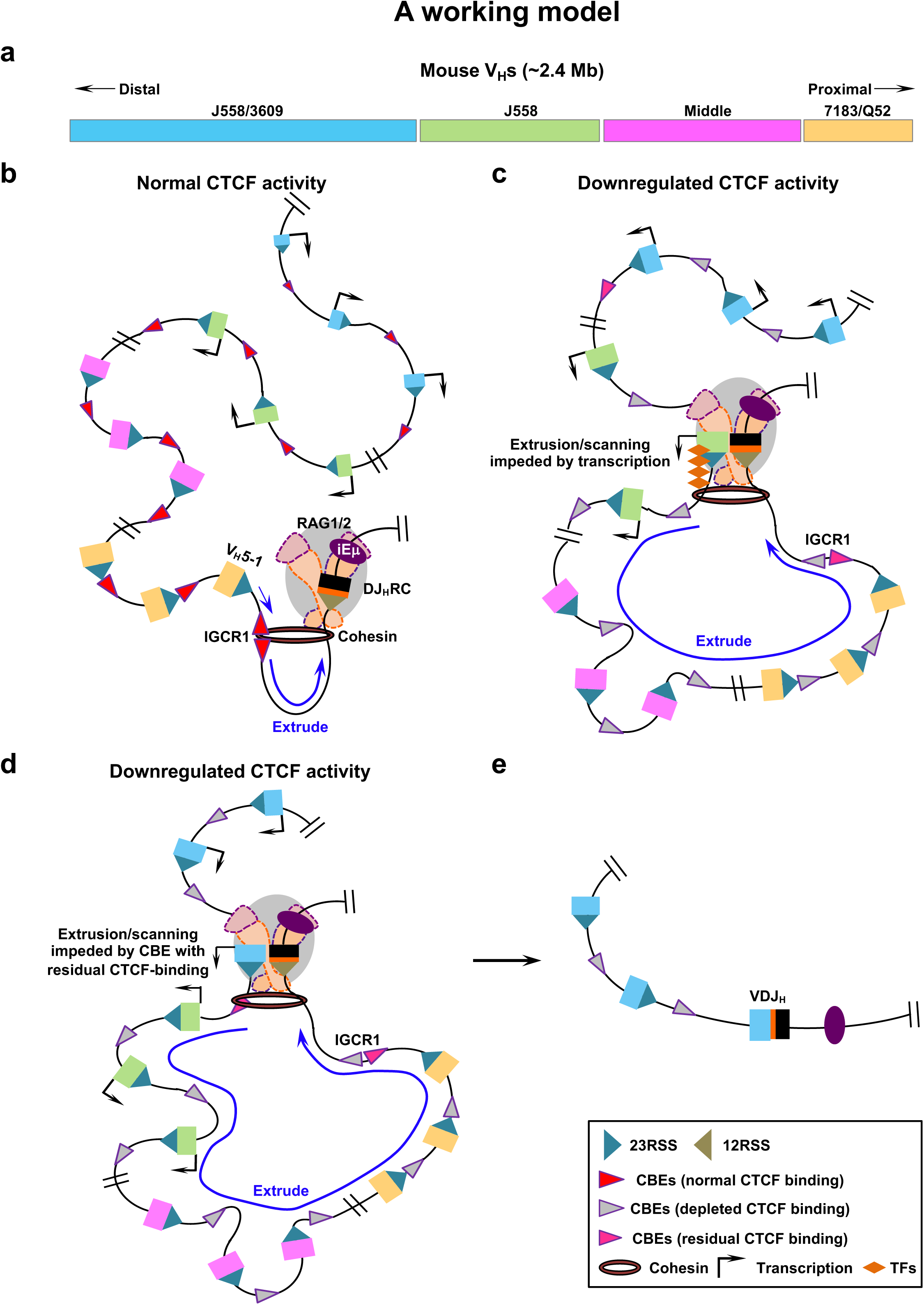
A working model for CTCF orchestrating long-range cohesin-driven V(D)J recombinational scanning. **a,** Diagram shows mouse V_H_ locus with 100-plus V_H_s in four domains^38, 39^, illustrated in different colors from J_H_-proximal to J_H_-distal. Besides harboring different V_H_ families, proximal domains are transcriptionally down-regulated and have CBEs immediately adjacent to V_H_-RSSs, whereas distal domains are transcriptionally active and have CBEs between but rarely adjacent to V_H_s. All V_H_ CBEs are oriented convergently with the IGCR1 upstream CBE1 and their RSSs convergently oriented with the DJ_H_ recombination center (RC)-RSS and must join by deletion. Details of V_H_ locus organization have been discussed^13, 15, 21–23, 38, 39^. **b,** With normal CBE-impediment activity (red arrowheads), RAG scanning is strongly impeded by CBE-based IGCR1 and non-CBE-based DJ_H_RC impediments^11, 15^, allowing scanning-based D-to-J_H_ rearrangement to proceed V_H_-to-DJ_H_ rearrangement^11^. Low-level scanning beyond IGCR1 allows proximal V_H_-to-DJ_H_ rearrangement in a minor subset of pro-B cells. Proximal V_H_-CBEs impedes further upstream scanning^15, 16^. **c, d,** Down-modulation of CTCF reduces CTCF-occupancy of CBEs (grey arrowheads) across *Igh*, but residual binding is retained at certain CBEs (pink arrowheads). Down-modulation of CTCF/CBE activity allows RC-bound RAG to scan various linear distances across the *Igh* in some cases the full-length. Upon reaching remaining scanning impediments, which could reflect transcription (dark arrows)^11, 37, 46^, transcription-factor (TF)-binding sites^15, 21–23, 38, 39^ (orange diamonds), and/or CBEs with residual CTCF binding, impeded scanning focuses RC-based RAG activity on sequences in the impeded region^11^ for V_H_-to-DJ_H_ joining^1^ (**e**). Down-modulation of CTCF binding activity is one manner for extending extrusion but this could also be achieved by circumventing CBE impediments, for example, through modulation of cohesin activity (See Text).

## METHODS

### Experimental procedures

No statistical methods were used to predetermine sample size. Experiments were not randomized and the investigators were not blinded to allocation during experiments and outcome assessment.

### Mice

Wild-type (WT) 129SV mice were purchased from Charles River Laboratories International. RAG2-deficient mice in 129SV background were generated^51^ and maintained in the Alt laboratory. The *RAG2^-/-^; Eμ-Bcl2^+^* mice were generated by crossing RAG2-deficient mice with *Eμ-Bcl2* transgenic mice^52^ that had been backcrossed to the 129SV background. All animal experiments were performed under protocols approved by the Institutional Animal Care and Use Committee of Boston Children’s Hospital.

### Bone marrow pro-B cell purification

Bone marrow (BM)-derived pro-B cells (B220^+^CD43^+^IgM^-^) were purified via FACS sorting as previously described^15, 40^ from 4-week-old WT 129SV mice. B220-positive BM pro-B cells were purified via anti-B220 biotin/streptavidin beads (Miltenyi, #130-049-501) from 4-week-old RAG2-deficient mice and were cultured in opti-MEM medium containing 10% (v/v) FBS plus IL-7/SCF for 4 days as previously described^14^.

### Cell lines

The parental *v-Abl*-kinase-transformed pro-B lines were derived by retroviral infection of BM pro-B cells derived from *RAG2^-/-^; Eμ-Bcl2^+^* mice with the pMSCV-v-Abl retrovirus, as described previously^25^. Infected cells were cultured in RPMI medium containing 15% (v/v) FBS for three months to obtain stably transformed *v-Abl* lines. The *RAG2^-/-^; Eμ-Bcl2^+^ v-Abl* pro-B lines were further targeted by Cas9/sgRNA approach^53^ to randomly introduce 1∼4 bp indels into the site ∼90 bp downstream of J_H_4-RSS-heptamer and ∼10 bp upstream of J_H_4-coding end (CE) bait primer on both *Igh* alleles. The *Igh* allele-specific barcodes permit the separation of the sequenced reads amplified from each allele with the identical bait primer from the same HTGT-V(D)J-seq libraries, thus, permitting in parallel examination of V(D)J recombination events that occur on each *Igh* allele under the same cellular context. HTGTS-V(D)J-seq analyses on RAG2-complemented above-mentioned *Igh*-barcoded *v-Abl* cells confirmed very comparable V(D)J rearrangement patterns and frequencies between *Igh* alleles, which were also very similar with those on germline *Igh* alleles in the parental *RAG2^-/-^; Eμ-Bcl2^+^ v-Abl* cells. The human colorectal carcinoma line HCT116 was purchased from American Type Culture Collection (ATCC, #CCL-247) and were cultured in McCoy’s 5A medium (ThermoFisher Scientific, #16600082) containing 10% (v/v) FBS.

### Generation of Rad21-degron and CTCF-degron *v-Abl* pro-B lines and their derivatives

The targeting constructs used for introducing the in-frame mAID sequences into mouse endogenous CTCF locus (pEN84, Addgene #86230) and for introducing the OsTir1-V5 expression cassette into endogenous *Rosa26* locus (pEN114, Addgene #92143) were described in a previously published study^9^. Using a similar strategy^9^, we generated the targeting construct for introducing the in-frame mAID sequences into mouse endogenous *Rad21* locus. Briefly, the 5’ and 3’ homology arms (1.5 kb each) flanking *Rad21* gene stop codon were PCR amplified from *RAG2^-/-^; Eμ-Bcl2^+^ v-Abl* pro-B cell genomic DNA (gDNA). The mAID-Clover cassette was PCR amplified from the plasmid pMK290 (Addgene, #72828). These fragments together with a puro selection cassette amplified from pEN84 were assembled in an order diagramed in Extended Data Fig. 1c using Gibson assembly kit (NEB, #E2611S). To promote the targeting efficiency, Cas9/sgRNAs were cloned by annealing pairs of oligos into pX330 (Addgene, #42230) following the protocol previously described^53^. The parental *RAG2^-/-^; Eμ-Bcl2^+^ v-Abl* pro-B cells were nucleofected with Rad21- or CTCF-targeting donors and their corresponding Cas9/sgRNAs via Lonza 4D Nucleofector as shown before^14, 15^, selected with 1 μg/ml Puromycin (Gibco, #A1113802) for 4 days, and subcloned by dilution. Candidate clones with desired gene modifications were screened by PCR and confirmed by Southern blot as outlined in Extended Data Fig. 1c and 4c. The confirmed clones were further processed to remove the puromycin selection cassette as previously described^9^. Similarly, the resultant clones were further targeted to introduce into both *Rosa26* alleles the OsTir1-V5 expression cassette followed by removal of the puromycin selection cassette as outlined in Extended Data Fig. 1d.

The resulting lines were referred to as the Rad21-degron or CTCF-degron *v-Abl* pro-B line and used in this study. To delete IGCR1, the Rad21-degron *v-Abl* pro-B line was targeted with Cas9/sgRNAs as previously described^14, 15^. To determine which *Igh* allele harbors IGCR1 deletion in the IGCR1^+/-^ Rad21-degron *v-Abl* clones, we developed and applied an efficient allele-specific assay based on the examination of Cas9/sgRNA-induced translocations via LAM-HTGTS^14, 54^. Briefly, we introduced the bait and prey Cas9/sgRNAs that, respectively, target the site located between J_H_4-RSS-heptamer and *Igh*-allele-specific barcodes and IGCR1 locale into the IGCR1^+/-^ Rad21-degron *v-Abl* clones, extracted the gDNA from the cells 3 days post the nucleofection, and performed LAM-HTGTS using J_H_4-CE bait primer. The bait-prey site translocation junctions derived from each of the two *Igh* alleles were separated through the barcodes, and the combined junction numbers from both strands that fall into bait and prey regions on each allele were plotted under a 1.5 Mb bin size. The relative numbers of prey junctions within IGCR1 locale indicate the presence or absence of IGCR1 on each allele, which can be further confirmed by HTGTS-V(D)J-seq analyses, as the rearrangements of proximal V_H_S are enhanced by IGCR1 deletion^14, 54^. Sequences of all sgRNAs and oligos used are listed in Supplementary Table 3.

### Treatment of Rad21-degron and CTCF-degron *v-Abl* pro-B lines with IAA

To deplete mAID-tagged Rad21 or CTCF protein in Rad21-degron or CTCF-degron *v-Abl* pro-B cells, the auxin analog, Indole-3-acetic acid (IAA, Sigma-Aldrich, #I3750-25G-A), was added in the medium at 500 μM from a 1000X stock that were prepared by dissolving IAA with DMSO. DMSO solvent was applied as the mock to non-treated (NT) cells. After 6 hours, the depletion efficiency of mAID-tagged proteins in IAA-treated cells compared to non-treated cells was examined by FACS. Both non-treated and IAA-treated cells were then treated by 3 μM STI-571 without or with IAA for 4 days to induce G1 arrest as previously described^11, 14, 54^. The cells were then collected and examined by FACS for protein depletion confirmation prior to various assays as described below.

### Cell viability assay

For time-course cell viability assay, Rad21-degron and CTCF-degron *v-Abl* pro-B cells were non-treated or treated with IAA for 6 hours and then subject to STI-571 treatment (set as 0 h). The viability of NT and +IAA cells at the following 4 days was determined by the percentage of viable lymphocyte population gated by FACS side (SSC) and forward (FSC) scatters out of the total cells. Average percentage ± s.d. of viable cells for each timepoint and for each condition was calculated from >3 biologically independent experiments and was normalized to the number of NT cells at 0 hours, which was set to 100%.

### G1 cell cycle stage analysis

For cell cycle analysis, the fluorescent, ubiquitination-based cell cycle indicator (Fucci) ^55^ cassette (pEN435, Addgene #92139) was stably introduced into the Rad21-degron and CTCF-degron *v-Abl* pro-B lines following the strategy as described previously^9^. The parental *v-Abl* cells without Fucci cassette, Rad21-degron and CTCF-degron *v-Abl* pro-B cells with Fucci cassette were non-treated or treated with IAA for 6 hours followed by STI-571 treatment (set as 0 h). The distribution of cells at G1 cell cycle stage (mCherry-hCdt1^+^; TagBFP-hGeminin^-^ population) at the following 4 days was determined by FACS using the parental *v-Abl* cells without Fucci cassette as the gating control. Average percentage ± s.d. of cells arrested in G1 stage for each timepoint and for each condition was calculated from >3 biologically independent experiments. The data collected at day 4 post STI-571 treatment are shown in Extended Data Fig. 1g, 4f.

### HTGTS-V(D)J-seq

For *Igh* V(D)J recombination analyses, we purified BM pro-B cells from WT 129SV mice as described above and introduced RAG2 into RAG2-deficient Rad21-degron and CTCF-degron *v-Abl* pro-B cells as well as their derivatives via the approach described previously^11^. HTGTS-V(D)J-seq libraries were prepared as previously described^11, 15, 40, 41^. Briefly, 2 μg of gDNA from sorted mouse BM pro-B cells or 30 μg of gDNA from G1-arrested RAG2-complemented parental *v-Abl* cells, Rad21-degron and CTCF-degron *v-Abl* cells without or with IAA treatment was sonicated and subjected to LAM-PCR using biotinylated J_H_4-CE bait primer^15^. Single-stranded LAM-PCR products were purified using Dynabeads MyONE C1 streptavidin beads (Life Technologies, #65002) and ligated to bridge adaptors. Adaptor-ligated products were amplified by nested PCR with indexed J_H_4 primers and the primer annealed to the adaptor. The PCR products were further tagged with Illumina sequencing adaptor sequences, size-selected via gel extraction and loaded onto an Illumina MiSeq machine for paired-end 250-bp or 300-bp sequencing. Primer sequences are listed in Supplementary Table 3.

### HTGTS-V(D)J-seq data processing and analyses

HTGTS-V(D)J-seq libraries were processed via the pipeline described previously^41^. *Igh* allele-specific sequencing reads were separated via barcodes and aligned to AJ851868/mm9 hybrid genome that was generated by combining all of the annotated *Igh* sequences of 129SV background (AJ851868) and the distal V_H_ sequences from the C57BL/6 background (mm9) starting from V_H_8-2 as previously described^15, 40, 56^. For statistical analyses, utilization data of V_H_ and D segments in G1-arrested CTCF-degron *v-Abl* pro-B cells was the sum up of the numbers obtained from both *Igh* alleles, each of which was normalized to 10,000 total recovered junctions as previously described^15^ and showed very similar utilization patterns. For comparisons, utilization data of V_H_ and D segments in BM pro-B cells was normalized to 20,000 total recovered junctions. We also extracted and re-analyzed the V(D)J recombination data obtained from 129SV BM pro-B cells in our prior publication^40^ and compared them with the newly generated data following the same normalization approach, which showed essentially same relative V_H_ and D utilization patterns. As the number of junctions used for normalization of Rad21 depletion experiments was greatly decreased comparing non-treated experiments due to the abrogation of nearly all V(D)J rearrangements by cohesin loss, we normalized the utilization data for Rad21 depletion experiments to the total aligned reads which include all bait primer-containing sequencing reads including V(D)J junctions for statistical analyses following the previously described strategy^14, 15^. In this regard, utilization data from G1-arrested Rad21-degron *v-Abl* pro-B cells with or without IAA treatment was the sum up of the numbers obtained from both *Igh* alleles, each of which was normalized to 110,000 total aligned reads. Utilization data from WT or IGCR1-deleted allele in G1-arrested IGCR1^+/-^ Rad21-degron *v-Abl* cells without or with IAA treatment was normalized to 110,000 total aligned reads. For each HTGTS-V(D)J-seq data plotted in figures and shown in Supplementary Table 1 and 2, average utilization frequencies ± s.d. were presented.

### 3C-HTGTS and data analyses

3C-HTGTS on short-term cultured RAG2-deficient BM pro-B cells, G1-arrested parental *v-Abl* cells, Rad21-degron and CTCF-degron *v-Abl* cells without or with IAA treatment, was performed as previously described^15^. Briefly, 10 million cells were crosslinked with 2% formaldehyde (Sigma-Aldrich, #F8775) for 10 min at room temperature and quenched with glycine at a final concentration of 125 mM. Cells were lysed on ice for 10 min followed by centrifugation to get nuclei. Nuclei were resuspended in NEB Cutsmart buffer for NlaIII (NEB, #R0125) digestion at 37 °C overnight, followed by T4 DNA ligase (Promega, #M1801) mediated ligation under dilute conditions at 16 °C overnight. Ligated products were treated with Proteinase K (Roche, #03115852001) and RNase A (Invitrogen, #8003089) followed by DNA purification to get the 3C templates. 3C-HTGTS library preparation follows the standard LAM-HTGTS library preparation procedures as previously described^15^. 3C-HTGTS libraries were sequenced via Illumina NextSeq550 using paired-end 150-bp sequencing kit or Miseq using paired-end 300-bp sequencing kit. Sequencing reads were processed as previously described^15^ and were aligned to AJ851868/mm9 hybrid genome as described above. Data were plotted for comparison after normalizing junction from each 3C-HTGTS library by random selection to the total number of chr12-wide junctions recovered from the smallest library in the set of comparing libraries. For G1-arrested Rad21-degron *v-Abl* pro-B cells with and without IAA treatment, 3C-HTGTS libraries using V_H_81X-CBE (Fig.1c, Extended Data Fig.2b) and iEμ (Fig.1c, Extended Data Fig.2c) as bait were normalized to 9,844 and 74,339 total junctions from chromosome 12, respectively. For G1-arrested IGCR1^-/-^ Rad21-degron *v-Abl* pro-B cells with and without IAA treatment, 3C-HTGTS libraries using iEμ as bait (Extended Data Fig.2d) were normalized to 74,339 total junctions from chromosome 12. For G1-arrested parental, non-treated and IAA-treated CTCF-degron *v-Abl* pro-B cells and RAG2-deficient BM pro-B cells, 3C-HTGTS libraries using RC/iEμ, RC/DQ52-J_H_1, and RC/J_H_4 as bait were normalized to 333,127 total junctions from chromosome 12 (Fig. 4b, Extended Data Fig.5c, 7b, c). Chromosomal interaction patterns were very comparable before and after normalization. The sequences of primers used for generating 3C-HTGTS libraries are listed in Supplementary Table 3.

### ChIP-seq

ChIP-seq was performed as previously described^11, 37^ on G1-arrested parental *v-Abl* cells, Rad21-degron and CTCF-degron *v-Abl* cells without or with IAA treatment. In brief, 20 million cells in fresh medium were crosslinked with 1% formaldehyde for 10 min at 37 °C, and quenched with glycine at final concentration of 125 mM. Crosslinked samples were collected and lysed in cell lysis buffer (5 mM PIPES pH 8.0, 85 mM KCl, 0.5% NP-40) followed by centrifugation to get the nuclei. Nuclei were lysed in nuclei lysis buffer (50mM Tris-HCl pH 8.1, 10mM EDTA, 1% SDS) and fragmented using Bioruptor at average size of 200-300 bp. Following the same procedure, fragmented chromatin from HCT116 cells were prepared and used as spike-in controls. Fragmented chromatin from G1-arrested cells with or without spike-in was precleared with 40 μl Dynabeads Protein A (ThermoFisher Scientific, #10002D) and then incubated with Rad21 (Abcam, #ab992) or CTCF (Millipore, #07-729) antibody in cold room overnight, followed by incubating with 40 μl Protein A beads for at least 2 hours. The immunoprecipitated DNA fragments were washed, eluted, de-crosslinked and purified for library preparation. ChIP-Seq libraries were prepared using NEBNext Ultra II DNA Library Prep Kit for Illumina (NEB, #E7645) and sequenced by paired-end 75-bp sequencing kit on Illumina NextSeq550.

### ChIP-seq analysis

We used bowtie2 to align reads to reference genome, used MACS2 to obtain the IP peak signal tracks after normalized to 1 million IP reads. Results were displayed in IGV. Rad21 ChIP-seq on Rad21-degron *v-Abl* pro-B cells with and without IAA treatment (Fig.1b, Extended Data Fig.2a) were performed with human HTC116 chromatin as spike-in^57^. For Rad21 ChIP-seq on parental and IAA-treated CTCF-degron *v-Abl* pro-B cells, one repeat was performed without human HCT116 chromatin as spike-in (Fig.2b, bottom panels), and two repeats were performed with human HCT116 chromatin as spike-in (Extended Data Fig.6a, bottom panels). Three repeats show very reproducible results. For CTCF ChIP-seq on parental and IAA-treated CTCF-degron *v-Abl* pro-B cells, all three repeats (Fig.2b, Extended Data Fig.6a, top panels) were performed without human HCT116 chromatin as spike-in. For ChIP-seq without human spike-in chromatin, reads were aligned to mouse AJ851868/mm9 hybrid genome and libraries were normalized to 1 million reads for display. For ChIP-seq with human spike-in chromatin, reads were aligned to the mixture genome of mouse (AJ851868/mm9) and human (hg19), and the mouse IP signal track was normalized to 1 million human IP reads, after adjusting the input spike-in chromatin ratio to be 1:1. To estimate and adjust for the actual spike-in chromatin ratio, we counted the number of reads aligned to human or mouse genome in the IP-corresponding input library, and calculated their ratio. We finally scaled the mouse IP signal track by multiplying its normalization factor = (1 million)/(human IP read number)/((mouse input read number)/(human input read number)) for display. For the samples that were processed with or without spike-in controls, we compared the results obtained from both ways and found little differences between them.

### GRO-seq and data analyses

GRO-seq libraries were prepared as described previously^11, 37^ on G1-arrested parental *v-Abl* cells, Rad21-degron and CTCF-degron *v-Abl* cells without or with IAA treatment. Briefly, 10 million cells were collected and permeabilized with the DEPC treated buffer (10 mM Tris-HCl pH 7.4, 300 mM sucrose, 10 mM KCl, 5 mM MgCl2, 1 mM EGTA, 0.05% Tween-20, 0.1% NP40 substitute, 0.5 mM DTT, protease inhibitors and RNase inhibitor). The permeabilized cells were resuspended in 100 μl of DEPC treated storage buffer (10 mM Tris-HCl pH 8.0, 25% (v/v) glycerol, 5 mM MgCl2, 0.1 mM EDTA and 5 mM DTT) followed by nuclear run-on with 100 μl 2X run-on mix (5 mM Tris-HCl pH 8.0, 2.5 mM MgCl2, 0.5 mM DTT, 150mM KCl, 0.5 mM ATP, 0.5 mM CTP, 0.5 mM GTP, 0.5 mM Br-UTP, RNase inhibitor, 1% Sarkosyl) at 37 °C for 5 min. Total RNA was extracted by Trizol and followed by hydrolyzation with NaOH at a final concentration of 0.2 N on ice for 18 min. After quenching with ice-cold Tris-HCl pH 6.8 at a final concentration of 0.55 M and exchanging buffer via Bio-Rad P30 columns, the total RNA was incubated with Br-dU antibody-conjugated beads (Santa Cruz, #sc-32323-ac) for 1 h. The enriched Br-dU labeled RNAs were incubated with RppH (NEB, #M0356S) and with T4 PNK (NEB, #M0201S) for hydroxyl repair, followed by ligating the 5′ and 3′ RNA adaptors. RT-PCR was performed after adaptor ligation to obtain cDNAs. Half of the cDNAs was subjected to library preparation by two rounds of PCR with barcoded primers. The second round of PCR products were purified by AMPure beads (Beckman Coulter, #A63880). GRO-seq libraries were sequenced via paired-end 75 bp sequencing on Illumina NextSeq550. For GRO-seq analysis, we used bowtie2 to align GRO-seq reads to AJ851868/mm9 hybrid genome, and run MACS2 to generate bigwig graph. To visualize the genome-wide RNA expression level, we run htseq-count to count read number on each gene, and made scatter plot after down-sampling each sample to 10 million reads mappable to any genes. We used DESeq2^58^ to call differentially expressed genes. Each experiment was repeated three times with multiple biologically independent clones.

### Quantification and statistical analysis

An unpaired, two-tailed Student’s *t* test was used to determine the statistical significance of differences between samples. At least three repeats were done for each statistical analysis. P values are calculated and shown in the figures as the follows: non-significant (NS): P > 0.05, *: P ≤ 0.05, **: P ≤ 0.01, and ***: P ≤ 0.001.

### Data availability

HTGTS-V(D)J-seq, 3C-HTGTS, ChIP-seq, and GRO-seq sequencing data reported in this study have been deposited in the GEO database under the accession number GSE142781. Specifically, HTGTS-V(D)J-seq data related to Fig. 1d, e, 3b-d, Extended Data Fig. 5b, and Supplementary Tables 1, 2 are deposited in the GEO database under the accession number GSE142777. 3C-HTGTS data related to Fig. 1c, 4b, and Extended Data Fig. 2b-d, 5c, 7b, c are deposited in the GEO database under the accession number GSE142778. ChIP-seq data related to Fig. 1b, 2b, and Extended Data Fig. 2a, 6a are deposited in the GEO database under the accession number GSE142779. GRO-seq data related to Fig. 2c, and Extended Data Fig. 3, 6b are deposited in the GEO database under the accession number GSE142780.

### Code availability

HTGTS-V(D)J-seq, 3C-HTGTS, ChIP-seq, and GRO-seq data were processed through the published pipelines as previously described^11, 15, 37^. Specifically, these pipelines are available at http://robinmeyers.github.io/transloc_pipeline/ (HTGTS pipeline), http://bowtie-bio.sourceforge.net/bowtie2/index.shtml (Bowtie2 v.2.2.8), https://sourceforge.net/projects/samtools/files/samtools/1.8/ (SAMtools v.1.8) and http://rseqc.sourceforge.net/# bam2wig-py (RSeQC tool v.2.6).

## REFERENCES

1. Alt, F. W., Zhang, Y., Meng, F.-L., Guo, C. & Schwer, B. Mechanisms of programmed DNA lesions and genomic instability in the immune system. Cell 152, 417–429 (2013).

2. Fudenberg, G. et al. Formation of Chromosomal Domains by Loop Extrusion. Cell Rep 15, 2038–2049 (2016).

3. Sanborn, A. L. et al. Chromatin extrusion explains key features of loop and domain formation in wild-type and engineered genomes. Proc. Natl. Acad. Sci. U.S.A. 112, E6456–65 (2015).

4. Rao, S. S. P. et al. Cohesin Loss Eliminates All Loop Domains. Cell 171, 305– 320.e24 (2017).

5. Haarhuis, J. H. I. et al. The Cohesin Release Factor WAPL Restricts Chromatin Loop Extension. Cell 169, 693–707.e14 (2017).

6. Schwarzer, W. et al. Two independent modes of chromatin organization revealed by cohesin removal. Nature 551, 51–56 (2017).

7. Wutz, G. et al. Topologically associating domains and chromatin loops depend on cohesin and are regulated by CTCF, WAPL, and PDS5 proteins. EMBO J. 36, 3573– 3599 (2017).

8. Vian, L. et al. The Energetics and Physiological Impact of Cohesin Extrusion. Cell 173, 1165–1178.e20 (2018).

9. Nora, E. P. et al. Targeted Degradation of CTCF Decouples Local Insulation of Chromosome Domains from Genomic Compartmentalization. Cell 169, 930–944.e22 (2017).

10. Rowley, M. J. & Corces, V. G. Organizational principles of 3D genome architecture. Nat. Rev. Genet. 19, 1–800 (2018).

11. Zhang, Y. et al. The fundamental role of chromatin loop extrusion in physiological V(D)J recombination. Nature 573, 600–604 (2019).

12. Guo, C. et al. CTCF-binding elements mediate control of V(D)J recombination. Nature 477, 424–430 (2011).

13. Lin, S. G., Guo, C., Su, A., Zhang, Y. & Alt, F. W. CTCF-binding elements 1 and 2 in the Igh intergenic control region cooperatively regulate V(D)J recombination. Proc. Natl. Acad. Sci. U.S.A. 112, 1815–1820 (2015).

14. Hu, J. et al. Chromosomal Loop Domains Direct the Recombination of Antigen Receptor Genes. Cell 163, 947–959 (2015).

15. Jain, S., Ba, Z., Zhang, Y., Dai, H.-Q. & Alt, F. W. CTCF-Binding Elements Mediate Accessibility of RAG Substrates During Chromatin Scanning. Cell 174, 102–116.e14 (2018).

16. Lin, S. G., Ba, Z., Alt, F. W. & Zhang, Y. RAG Chromatin Scanning During V(D)J Recombination and Chromatin Loop Extrusion are Related Processes. Adv. Immunol. 139, 93–135 (2018).

17. Degner, S. C. et al. CCCTC-binding factor (CTCF) and cohesin influence the genomic architecture of the Igh locus and antisense transcription in pro-B cells. Proc. Natl. Acad. Sci. U.S.A. 108, 9566–9571 (2011).

18. Lucas, J. S., Bossen, C. & Murre, C. Transcription and recombination factories: common features? Curr. Opin. Cell Biol. 23, 318–324 (2011).

19. Lucas, J. S., Zhang, Y., Dudko, O. K. & Murre, C. 3D trajectories adopted by coding and regulatory DNA elements: first-passage times for genomic interactions. Cell 158, 339–352 (2014).

20. Khanna, N., Zhang, Y., Lucas, J. S., Dudko, O. K. & Murre, C. Chromosome dynamics near the sol-gel phase transition dictate the timing of remote genomic interactions. Nat Commun 10, 2771 (2019).

21. Bossen, C., Mansson, R. & Murre, C. Chromatin topology and the regulation of antigen receptor assembly. Annu. Rev. Immunol. 30, 337–356 (2012).

22. Proudhon, C., Hao, B., Raviram, R., Chaumeil, J. & Skok, J. A. Long-Range Regulation of V(D)J Recombination. Adv. Immunol. 128, 123–182 (2015).

23. Ebert, A., Hill, L. & Busslinger, M. Spatial Regulation of V-(D)J Recombination at Antigen Receptor Loci. Adv. Immunol. 128, 93–121 (2015).

24. Muljo, S. A. & Schlissel, M. S. A small molecule Abl kinase inhibitor induces differentiation of Abelson virus-transformed pre-B cell lines. Nat. Immunol. 4, 31–37 (2003).

25. Bredemeyer, A. L. et al. ATM stabilizes DNA double-strand-break complexes during V(D)J recombination. Nature 442, 466–470 (2006).

26. Nishimura, K., Fukagawa, T., Takisawa, H., Kakimoto, T. & Kanemaki, M. An auxin-based degron system for the rapid depletion of proteins in nonplant cells. Nat. Methods 6, 917–922 (2009).

27. Natsume, T., Kiyomitsu, T., Saga, Y. & Kanemaki, M. T. Rapid Protein Depletion in Human Cells by Auxin-Inducible Degron Tagging with Short Homology Donors. Cell Rep 15, 210–218 (2016).

28. Yatskevich, S., Rhodes, J. & Nasmyth, K. Organization of Chromosomal DNA by SMC Complexes. Annu. Rev. Genet. (2019). doi:10.1146/annurev-genet-112618-043633

29. Hassler, M., Shaltiel, I. A. & Haering, C. H. Towards a Unified Model of SMC Complex Function. Curr. Biol. 28, R1266–R1281 (2018).

30. Uhlmann, F. SMC complexes: from DNA to chromosomes. Nat. Rev. Mol. Cell Biol. 17, 399–412 (2016).

31. Haarhuis, J. H. I., Elbatsh, A. M. O. & Rowland, B. D. Cohesin and its regulation: on the logic of X-shaped chromosomes. Dev. Cell 31, 7–18 (2014).

32. Peters, J.-M., Tedeschi, A. & Schmitz, J. The cohesin complex and its roles in chromosome biology. Genes Dev. 22, 3089–3114 (2008).

33. Wu, N. & Yu, H. The Smc complexes in DNA damage response. Cell Biosci 2, 5 (2012).

34. Davidson, I. F. et al. DNA loop extrusion by human cohesin. Science (2019). doi:10.1126/science.aaz3418

35. Kim, Y., Shi, Z., Zhang, H., Finkelstein, I. J. & Yu, H. Human cohesin compacts DNA by loop extrusion. Science 366, 1345–1349 (2019).

36. Zhao, L. et al. Orientation-specific RAG activity in chromosomal loop domains contributes to Tcrd V(D)J recombination during T cell development. J. Exp. Med. 213, 1921–1936 (2016).

37. Zhang, X. et al. Fundamental roles of chromatin loop extrusion in antibody class switching. Nature 575, 385–389 (2019).

38. Choi, N. M. et al. Deep sequencing of the murine IgH repertoire reveals complex regulation of nonrandom V gene rearrangement frequencies. J. Immunol. 191, 2393– 2402 (2013).

39. Bolland, D. J. et al. Two Mutually Exclusive Local Chromatin States Drive Efficient V(D)J Recombination. Cell Rep 15, 2475–2487 (2016).

40. Lin, S. G. et al. Highly sensitive and unbiased approach for elucidating antibody repertoires. Proc. Natl. Acad. Sci. U.S.A. 113, 7846–7851 (2016).

41. Hu, J. et al. Detecting DNA double-stranded breaks in mammalian genomes by linear amplification-mediated high-throughput genome-wide translocation sequencing. Nat Protoc 11, 853–871 (2016).

42. Aiden, E. L. & Casellas, R. Somatic Rearrangement in B Cells: It’s (Mostly) Nuclear Physics. Cell 162, 708–711 (2015).

43. Nakahashi, H. et al. A genome-wide map of CTCF multivalency redefines the CTCF code. Cell Rep 3, 1678–1689 (2013).

44. Canzio, D. et al. Antisense lncRNA Transcription Mediates DNA Demethylation to Drive Stochastic Protocadherin α Promoter Choice. Cell (2019). doi:10.1016/j.cell.2019.03.008

45. Ghirlando, R. & Felsenfeld, G. CTCF: making the right connections. Genes Dev. 30, 881–891 (2016).

46. Hsieh, T.-H. S. et al. Resolving the 3D landscape of transcription-linked mammalian chromatin folding. bioRxiv 176, 638775 (2019).

47. Xiang, Y., Park, S.-K. & Garrard, W. T. A major deletion in the Vκ-Jκ intervening region results in hyperelevated transcription of proximal Vκ genes and a severely restricted repertoire. J. Immunol. 193, 3746–3754 (2014).

48. Ribeiro de Almeida, C. et al. The DNA-binding protein CTCF limits proximal Vκ recombination and restricts κ enhancer interactions to the immunoglobulin κ light chain locus. Immunity 35, 501–513 (2011).

49. Seitan, V. C. et al. A role for cohesin in T-cell-receptor rearrangement and thymocyte differentiation. Nature 476, 467–471 (2011).

50. Chen, S. & Krangel, M. S. Diversification of the TCR β Locus Vβ Repertoire by CTCF. Immunohorizons 2, 377–383 (2018).

## References

51. Shinkai, Y. et al. RAG-2-deficient mice lack mature lymphocytes owing to inability to initiate V(D)J rearrangement. Cell 68, 855–867 (1992).

52. Strasser, A. et al. Enforced BCL2 expression in B-lymphoid cells prolongs antibody responses and elicits autoimmune disease. Proc. Natl. Acad. Sci. U.S.A. 88, 8661– 8665 (1991).

53. Cong, L. et al. Multiplex genome engineering using CRISPR/Cas systems. Science 339, 819–823 (2013).

54. Frock, R. L. et al. Genome-wide detection of DNA double-stranded breaks induced by engineered nucleases. Nat. Biotechnol. 33, 179–186 (2015).

55. Sakaue-Sawano, A. et al. Visualizing spatiotemporal dynamics of multicellular cell-cycle progression. Cell 132, 487–498 (2008).

56. Medvedovic, J. et al. Flexible long-range loops in the VH gene region of the Igh locus facilitate the generation of a diverse antibody repertoire. Immunity 39, 229–244 (2013).

57. Orlando, D. A. et al. Quantitative ChIP-Seq normalization reveals global modulation of the epigenome. Cell Rep 9, 1163–1170 (2014).

58. Love, M. I., Huber, W. & Anders, S. Moderated estimation of fold change and dispersion for RNA-seq data with DESeq2. Genome Biol. 15, 550 (2014).

